# Sonic hedgehog signaling in astrocytes mediates cell type-specific synaptic organization

**DOI:** 10.1101/537860

**Authors:** Steven Hill, Andrew S. Blaeser, Austin A. Coley, Yajun Xie, Katherine A. Shepard, Corey C. Harwell, Wen-Jun Gao, A. Denise R. Garcia

## Abstract

Astrocytes have emerged as integral partners with neurons in regulating synapse formation and function, but the mechanisms that mediate these interactions are not well understood. Here, we show that Sonic hedgehog (Shh) signaling in mature astrocytes is required for establishing structural organization and remodeling of cortical synapses in a cell type-specific manner. In the postnatal cortex, Shh signaling is active in a subpopulation of mature astrocytes localized primarily in deep cortical layers. Selective disruption of Shh signaling in astrocytes produces a dramatic increase in synapse number specifically on layer V apical dendrites that emerges during adolescence and persists into adulthood. Dynamic turnover of dendritic spines is impaired in mutant mice and is accompanied by an increase in neuronal excitability and a reduction of the glial-specific, inward-rectifying K^+^ channel Kir4.1. These data identify a critical role for Shh signaling in astrocyte-mediated modulation of neuronal activity required for sculpting synapses.

## INTRODUCTION

The organization of synapses into the appropriate number and distribution occurs through a process of robust synapse addition followed by a period of refinement during which excess synapses are eliminated. Failure to establish or maintain appropriate synaptic organization is a hallmark of many neurodevelopmental disorders^1^. Considerable evidence now shows that, together with neurons, astrocytes are critical regulators of synaptic connectivity and function^2–4^. Astrocytes interact intimately with synapses to regulate their formation, maturation, and function, and a growing number of astrocyte-secreted proteins that directly mediate synapse formation and elimination have been identified^5–9^. In addition, astrocytes regulate concentrations of K^+^ and glutamate in the extracellular space, thereby modulating neuronal activity^10^. Nevertheless, despite the remarkable progress in our understanding of the essential role for astrocytes in regulating synaptic formation and function, the underlying signaling programs mediating astrocyte-dependent regulation of synapse organization remain poorly understood.

The molecular signaling pathway Sonic hedgehog (Shh) governs a broad array of neurodevelopmental processes in the vertebrate embryo, including morphogenesis, cell proliferation and specification, and axon pathfinding^11,12^. However, Shh activity persists in multiple cell populations in the postnatal and adult CNS, including progenitor cells, as well as in differentiated neurons and astrocytes^13–17^, where novel and unexpected roles for Shh activity are emerging^18^. Following injury, Shh has been shown to mitigate inflammation^19,20^, and in the cerebellum, Shh derived from Purkinje neurons instructs phenotypic properties of mature Bergmann glia^21^. In the postnatal cortex, Shh is required for establishing local circuits between two distinct projection neuron populations^16^. Shh produced by layer V neurons guides the formation of synaptic connections to its layer II/III presynaptic partners, which transduce the Shh signal through non-canonical, Gli-independent mechanisms. We have previously shown that Shh signaling is also active in a discrete subpopulation of cortical astrocytes^17^, suggesting that Shh signaling mediates both homotypic and heterotypic cellular interactions. Astrocytes engaging in Shh activity are identified by expression of Gli1, a transcriptional effector of canonical Shh signaling^11^. Whether Shh signaling in cortical astrocytes plays a role in synaptic organization of neurons is not known.

In this study, we examined the organization and dynamics of dendritic spines on cortical neurons following selective disruption of Shh signaling in astrocytes. Dendritic spines are the structural hosts of most excitatory synapses and play an important role in the organization and function of neural circuits. We show that deep layer neurons exhibit long-lasting aberrations in the density and turnover of dendritic spines that emerge during postnatal development following selective disruption of Shh signaling in astrocytes. These perturbations in synaptic organization are not observed in upper layer neurons where Gli1 astrocytes are relatively sparse. Chronic *in vivo* imaging of dendritic spines reveals that mutant mice exhibit lower rates of spine turnover, suggesting impaired structural plasticity. In addition, these mice show a pronounced deficit in expression of the glial-specific inward-rectifying K^+^ channel, Kir4.1, as well as an increase in excitability of cortical neurons. Taken together, these data demonstrate that astrocytes act as key modulators of neural activity and structure during postnatal development in a Shh-dependent manner and further establish Shh signaling as a fundamental mediator of synaptic connectivity.

## RESULTS

### Gli1 astrocytes exhibit a distinct laminar distribution in the adult cortex

In the mature cortex, a subpopulation of astrocytes express the transcription factor Gli1, indicating active Shh signaling^17^. Notably, the distribution of Gli1 astrocytes throughout the cortex is non-uniform, showing a laminar-specific pattern (**Figure 1**). We analyzed the laminar distribution of Gli1 astrocytes in adult *Gli1*^*CreER*/+^;Ai14 mice, in which tamoxifen administration promotes Cre-mediated recombination of the fluorescent tdTomato reporter protein, permanently marking Gli1-expressing cells. Adult *Gli1*^*CreER*/+^;Ai14 mice received tamoxifen over three days, and their brains were analyzed two – three weeks later. In our previous study, we observed weak expression of βGal reporter protein in *Gli1*^*CreER*/+^*;R26^lacZ/lacZ^* mice at time points earlier than 4 weeks after tamoxifen^17^. In contrast, expression of the tdTomato reporter protein in the Ai14 reporter line shows robust expression as early as two weeks, and a comparable number, distribution and intensity of reporter-positive cells between two and four weeks^20^. For these studies, we therefore used a two – three week chase after tamoxifen. The vast majority of marked cells were observed within layers IV and V, with few marked cells observed in layers II/III or VI (**Figure 1**). We performed immunostaining with the pan-astrocytic marker S100β and quantified the fraction of marked astrocytes in cortical layers (**Figure 1**). This analysis showed that 44% of astrocytes in layer IV express Gli1 (**Figure 1**), while 36% of astrocytes in layer V express Gli1. Interestingly, the distribution of Gli1 astrocytes in layer V was not homogenous, showing an enrichment of marked cells in layer Vb, and a distinctive paucity of marked cells in layer Va (**Figure 1**), consistent with the localization of Shh producing neurons in layer Vb^16,17^. We analyzed the fraction of Gli1 astrocytes in each sublayer of layer V and found that 75% of marked cells were found in layer Vb whereas only 25% were found in layer Va. Layers II/III and VI showed the lowest fractions of marked astrocytes, with only 11% and 22% of S100β cells expressing tomato, respectively. These data demonstrate that Gli1 astrocytes are preferentially localized in deep cortical layers and suggest their interactions with local neurons may differ from that of Gli1-negative astrocytes in superficial layers.

**Figure 1:**
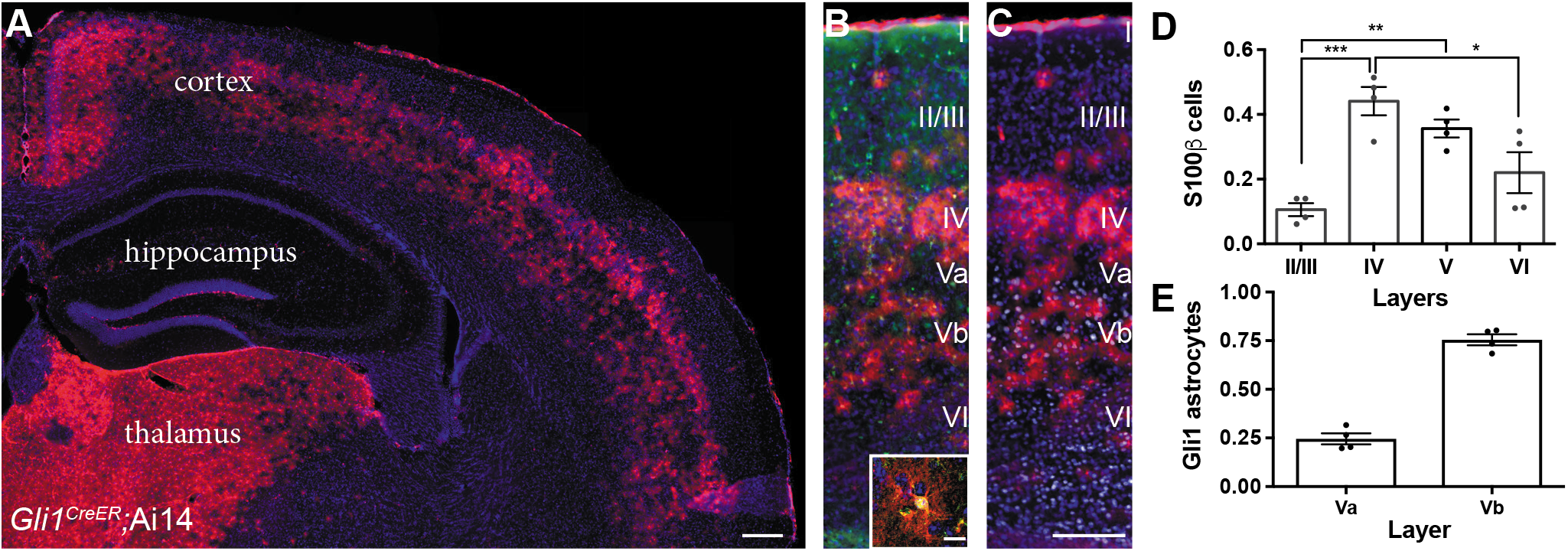
Gli1 astrocytes are distributed in a laminar-specific manner. **(A)** Tamoxifen-induced reporter expression in *Gli1^CreER^;*Ai14 mice reveals a laminar distribution of cells undergoing active Shh signaling (red). Scale bar, 250 μm. **(B)** Gli1+ cells (red) are a subset of all astrocytes (S100β, green, see inset; scale bar, 12.5 μm). **(C)** Layer analysis of Gli1 astrocytes (red) with DAPI and the deep layer marker Ctip2 (gray) illustrate their layer-specific organization. Scale bar, 125 μm. **(D)** The fraction of astrocytes expressing Gli1 in each cortical layer. Data points indicate individual animals, bars represent mean ± SEM. Statistical signficance was assessed by one-way ANOVA with Tukey’s post hoc test for multiple comparisons. *, *p* < 0.05; **, *p* < 0.01; ***, *p* < 0.01. **(E)** The fraction of Gli1 astrocytes in sublayers Va and Vb. Bar graphs represent mean value ± SEM. *n* = 4 animals.

### Shh signaling is required for cell type-specific synaptic connectivity

Transduction of canonical Shh signaling begins with binding of Shh to its transmembrane receptor, Patched1 (Ptch), relieving inhibition of a second, obligatory coreceptor Smoothened (Smo). To investigate the role of Shh signaling in astrocyte function, we performed conditional knockout of Smo selectively in astrocytes using GFAP-Cre transgenic mice in which Cre expression is regulated by the full-length mouse GFAP promoter^22^ (mGFAP Smo CKO). Because mGFAP-mediated Cre recombination targets astrocyte progenitors expressing GFAP which are present at birth, this is an effective tool for targeting the cortical astrocyte population for selective gene deletion. We validated the Cre-mediated recombination pattern in mGFAP-Cre mice by crossing them with Ai14 reporter mice (*mGFAP-Cre;Ai14*), that express the tdTomato reporter protein. Although individual floxed alleles possess distinct recombination efficiencies, owing to specific characteristics of each allele including distance between loxP sites or their accessibility due to chromatin structure^23^, analysis of recombination in a reporter line provides a useful approximation of the expression pattern of a given Cre driver. We performed single cell analysis of the identity of recombined cells. The vast majority of tomato-positive cells showed a bushy morphology, typical of protoplasmic astrocytes (**Figure 2 – figure supplement 1**). Single cell analysis of double staining with S100β showed that nearly 70% were co-labeled, identifying them as astrocytes. We also identified a small fraction (6%) of tomato cells as oligodendrocytes. A small fraction (9%) of tomato cells were co-labeled with the neuronal marker, NeuN, but this represented only 1.3% of cortical neurons (**Figure 2 – figure supplement 1**). Importantly, although a minor fraction of recombined cells was identified as neurons or oligodendrocytes, nearly all cortical astrocytes analyzed expressed the tdTomato reporter protein (95%; **Figure 2 – figure supplement 1**), suggesting effective targeting of the cortical astrocyte population using this mGFAP-Cre driver.

We performed qPCR on whole cortex from mGFAP Smo CKO mice and littermate controls and found a 50% reduction in the number of Smo transcripts (**Figure 2 - figure supplement 2**). As Smo is also expressed in neurons^16^, the remaining Smo transcripts are likely due to neurons that do not undergo Cre-mediated recombination in these mice. Importantly, mGFAP Smo CKO mice show a nearly complete loss of Gli1 activity, with no difference in the number of astrocytes in the mature cortex, demonstrating effective interruption of canonical Shh signaling^17^. These data demonstrate that this mGFAP-Cre driver is an effective tool for selective deletion of Smo in cortical astrocytes, and that canonical, Gli-mediated Shh activity is effectively abolished

During postnatal development, Shh is required for establishing synaptic connectivity between layer V neurons and their presynaptic partners in layer II/III, which is mediated by Gli-independent, non-canonical signaling^16^. To examine whether canonical, Gli-mediated Shh signaling in astrocytes plays a role in establishing synaptic connectivity of cortical neurons, we evaluated spine density of apical dendrites on layer V neurons in the somatosensory cortex of mGFAP Smo CKO mice across postnatal development. Dendritic spines receive the vast majority of excitatory input, thereby serving as a useful readout of synaptic connectivity^24^. To visualize neurons for reliable identification and tracing, mGFAP Smo CKO mice were crossed with Thy1-GFPm transgenic mice, which sparsely express GFP in a subset of layer V cortical neurons^25^. Dendritic spines undergo a period of dynamic reorganization during postnatal development during which there is an initial overproduction of spines over the first 2-3 weeks of postnatal development followed by a period of synaptic pruning that refines the precise connectivity of cortical circuits^26^. At P14, spine density was comparable between mGFAP Smo CKO mice and wild type (WT) littermate controls, suggesting that early synaptogenesis does not require astrocytic Shh signaling (0.58 ± 0.03 spines/μm and 0.56 ± 0.01 spines/μm, WT [*n* = 7 cells] and mGFAP Smo CKO [*n* = 6 cells], respectively, *p* = 0.64, 2 animals per genotype). Between P14 and P21, spine density remained stable in WT, but showed a dramatic increase at P28 (**Figure 2**). This period of synapse addition was followed by a steady reduction in spine density at P42 and into adulthood (≥P90), reflecting the developmental elimination of spines. In contrast, mGFAP Smo CKO mice showed an accelerated timeline of spine addition in which spine density increased dramatically between P14 and P21 (**Figure 2**). Spine density was significantly higher at P21 in mGFAP Smo CKO mice compared to controls (0.53 ± 0.05 spines/μm and 0.76 ± 0.06 spines/μm, WT [*n* = 9 cells] and mGFAP Smo CKO [*n* = 9 cells], respectively, *p* = 0.008, 3 animals per genotype). Although there was a modest reduction in spine density as animals matured, spine density remained elevated in adult mGFAP Smo CKO mice compared to WT controls (0.36 ± 0.02 spines/μm and 0.60 ± 0.02 spines/μm in WT [*n* = 24 cells] and mGFAP Smo CKO [*n* = 23 cells], respectively, *p* < 0.0001, 4 animals per genotype; **Figure 2**). Thus, whereas WT mice experience a 49% reduction in spine density from the peak of spine density at P28 to adulthood, mGFAP Smo CKO mice exhibit a 22% reduction from their peak at P21 to adulthood. Interestingly, we find no difference in basal dendrite spine density in mGFAP Smo CKO mice (0.60 ± 0.04 spines/um and 0.62 ± 0.04 spines/μm in WT [*n* = 10 cells] and mGFAP Smo CKO [*n* = 10 cells], respectively, *p* = 0.71, 3 animals per genotype; **Figure 2 – figure supplement 3**). This is in contrast to embryonic deletion of Shh from neurons using the Emx-Cre driver, which produces a reduction in spine density of basal dendrites on layer V neurons^16^. These data suggest that while the early stages of synapse formation proceed independently of astrocytic Shh signaling, the maturing cortical circuit requires intact Shh activity in astrocytes for the developmental pruning of excess synapses necessary to achieve its mature organization.

**Figure 2:**
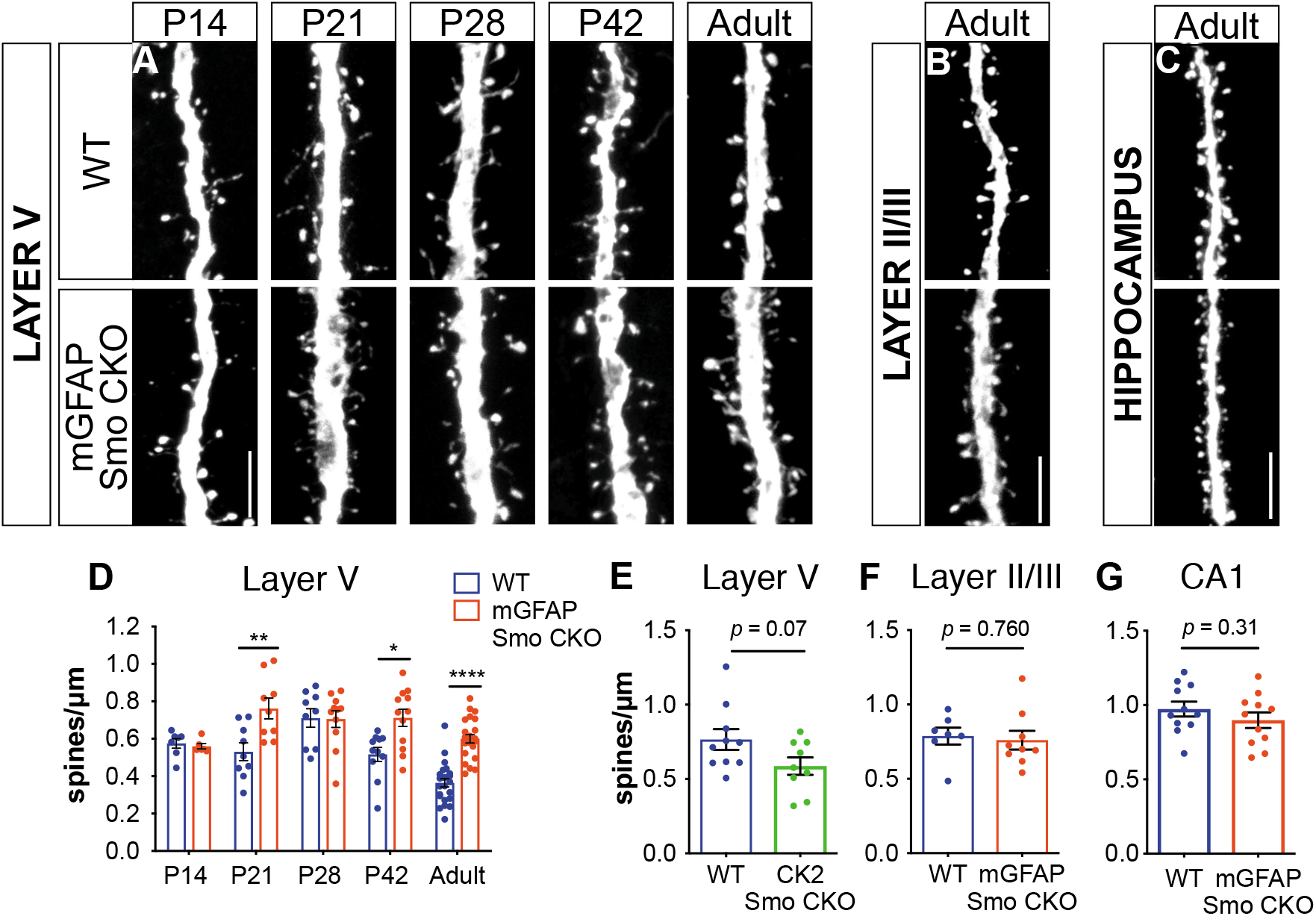
Loss of Shh signaling in astrocytes results in increased spine density on layer V neurons. **(A)** GFP immunostaining of representative apical dendritic segments from layer V neurons in WT and mGfap Smo CKO mice at all ages and regions analyzed. Scale bar, 5 *μ*m. **(D)** Spine density of apical dendrites from layer V neurons across various postnatal ages and in adult mice. Data points represent individual cells, bars represent mean ± SEM. Statistical analysis performed by two-way ANOVA with Tukey’s post-hoc test for multiple comparisons. *, *p* < 0.05; **, *p* < 0.01; ****, *p* < 0.0001. Significance is stated as mGFAP Smo CKO compared to WT at a given age. **(E)** Spine density of apical dendrites from layer V neurons in adult CamKlla Smo CKO mice. **(F-G)** Spine density of apical dendrites from layer II/III (F) and CA1 pyramidal neurons (G) in WT and mGfap Smo CKO mice. Data points represent individual cells, bars represent mean ± SEM. Statistical signficance assessed by Student’s t-test. *n* ≥ 3 animals per genotype for all ages except P14, where *n* = 2 animals per genotype.

To confirm that elevated spine density was due specifically to a loss of astrocytic Shh signaling, we interrogated the spine density of pyramidal cells in two different regions where Gli1 astrocytes are relatively sparse (see **Figure 1**). Pyramidal neurons in layer II/III of WT mice showed a higher spine density than layer V neurons (0.79 ± 0.06 spines/μm, *n* = 7 cells, 4 animals) consistent with previous studies^27^. However, this was not significantly different from the spine density observed in mGFAP Smo CKO mice (0.76 ± 0.06 spines/μm, *p* = 0.76, *n* = 9 cells, 5 animals; Figure 2). We also analyzed pyramidal neurons in the hippocampus. Although the dentate gyrus of the hippocampus harbors a population of Gli1-expressing adult neural progenitor cells, mature, differentiated astrocytes expressing Gli1 are relatively sparse^14,28^ (Figure 1). The spine density of CA1 pyramidal neurons in adult mice was higher than in cortical neurons, consistent with previous studies^29,30^. However, there was no significant difference in spine density between mGFAP Smo CKO mice and WT controls (0.97 ± 0.05 spines/μm and 0.90 ± 0.05 spines/μm in WT [*n* = 11 cells] and mGFAP Smo CKO [*n* = 11 cells], respectively, *p* = 0.32, 3 animals per genotype; **Figure 2**). Our analysis of the recombination pattern of the mGFAP-Cre driver showed that some cortical neurons undergo recombination (**Figure 2 – figure supplement 1**). To rule out the possibility that Smo deletion in a small population of neurons is responsible for the elevated spine density of layer V neurons, we deleted Smo in excitatory neurons using the CamKIIa-Cre driver (CK2 Smo CKO), and evaluated spine density of layer V neurons in adult mice. We found no significant difference in spine density between CK2 Smo CKO mice and littermate controls (0.77 ± 0.07 spines/μm and 0.59 ± 0.06 spines/μm in WT [*n* = 10 cells] and CK2 Smo CKO [*n* = 9 cells], respectively, *p* = 0.07, 3 animals per genotype; **Figure 2**). Taken together, these data suggest that Shh activity in astrocytes is necessary for the developmental reorganization of dendritic spines specifically on layer V cortical neurons. Notably, we observed elevated spine density along the apical dendrites of layer V neurons, which traverse layer II/III, where few Gli1 astrocytes are found. This suggests that the modification of synapse organization mediated by Gli1 astrocytes does not occur through direct astrocyte-synapse interactions.

### Synapses are more stable in mGFAP Smo CKO mice

Dendritic spines are dynamic structures that undergo rapid formation and elimination during postnatal cortical development. As cortical circuits mature, the fraction of spines undergoing dynamic turnover declines concomitant with an increase in spine stability^26,27^. The dynamic turnover of spines has long been considered a structural correlate of synaptic plasticity and can be regulated in an activity-dependent manner^31–34^. To evaluate the role of astrocytic Shh signaling in mediating spine dynamics, we performed repeated *in vivo* imaging of the apical tufts of layer V neurons in the somatosensory cortex through a cranial window^35^. We confirmed the layer V identity of imaged neurons by following individual dendritic segments to their soma and created 3D reconstructions of imaged neurons. Only dendrites with soma in layer V, typically 500 μm – 600 μm from the surface of the brain, were analyzed. In Thy1-GFPm mice, expression of GFP is developmentally regulated such that at P14, the intensity of GFP expression and the density of fluorescently labeled neurons within the 3 mm cranial window is insufficient for reliable imaging, as has been previously reported^27^. However by P17, GFP expression was sufficiently dense and bright to enable reliable imaging. We analyzed the fraction of spines undergoing dynamic turnover over 2 days in young mice at P17-P21 and P28-P32. In WT mice, the turnover ratio was 0.14 at P17-P21 (*n* = 5 mice) and 0.10 in P28-P32 mice (*n* = 3 mice; **Figure 3**). In mGFAP Smo CKO mice however, the turnover ratio was lower at P17-P21 than in WT mice, and remained stable at P28-P32 (0.11 and 0.10, respectively, *n* = 4 mice; **Figure 3**). This suggests that dendritic spines in mGFAP Smo CKO mice stabilize earlier than those in WT mice. Consistent with this, we found that the fraction of filopodia was lower in juvenile mGFAP Smo CKO mice compared to WT controls (**Figure 3**). Filopodia are structural precursors of dendritic spines and exhibit higher rates of dynamic turnover than mature spines^36,37^. Accordingly, the density of filopodia declines as the cortex matures^26,38^. Indeed, the fraction of all protrusions in juvenile WT mice (*n* = 7 mice) identified as filopodia was 25%, but that fraction declined significantly to 15% in adults (*n* = 16 mice, *p* = 0.0039; **Figure 3**). In contrast, we did not observe this age-dependent decline in mGFAP Smo CKO mice (17% and 12% in juvenile [*n* = 10 mice] and adult [*n* = 7 mice] mGFAP Smo CKO mice, respectively, *p* = 0.398). The filopodial fraction in juvenile mGFAP Smo CKO mice was already significantly below WT levels and comparable to adult WT levels, suggesting that nascent spines undergo accelerated stabilization in the absence of astrocyte-mediated Shh signaling.

**Figure 3:**
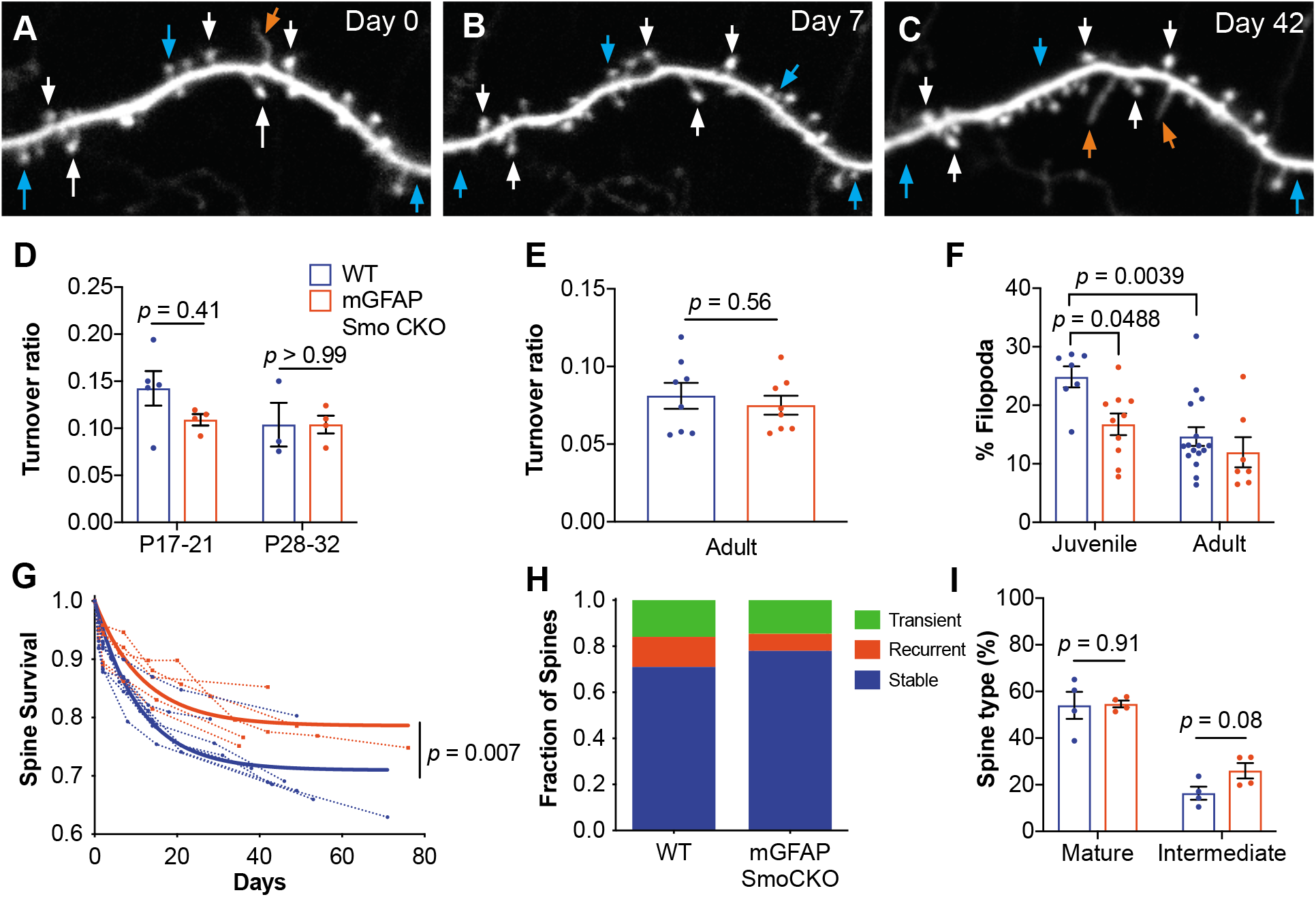
Loss of astrocytic Shh signaling impairs structural plasticity. **(A-C)** Example dendrite segment imaged repeatedly over 6 weeks. Day 0 indicates the first day of imaging, subsequent imaging days indicated. White arrows, stable spines; cyan arrows, transient spines; orange arrows, filopodia. **(D)** Turnover ratios in WT (*n* = 3-5 animals per age group) and mGFAP Smo CKO juvenile mice (*n* = 4 animals per age group) analyzed over 2 days. **(E)** Turnover ratios in WT (*n* = 8 animals) and mGFAP Smo CKO (*n* = 8 animals) adult mice analyzed over 7 days. Statistical analysis by two-way ANOVA, Tukey’s post hoc test (D) and unpaired Student’s t-test (E). **(F)** Fraction of protrusions identified as filopodia in juvenile (WT, *n* = 7 animals; mGFAP Smo CKO, *n* = 10 animals) and adult mice (WT, *n* = 16: mGFAP Smo CKO, *n* = 7 animals). Statistics by two-way ANOVA, Tukey’s post hoc test. **(G)** Comparison of spine survival curves for WT and mGFAP Smo CKO neurons. Each dashed curve represents the curve from an individual mouse (WT, *n* = 7 animals; mGFAP Smo CKO, *n* = 5 animals). Solid curves represent best-fits to exponential decay models. **(H)** The fraction of spines belonging to the stable, recurrent, and transient populations. Statistical significance was assessed by Student’s t-test for each class (stable, recurrent, or transient; *n* = 7 and 8 animals, WT and mGFAP Smo CKO, respectively). Stable, *p* = 0.0485; Recurrent, *p* = 0.1023; Transient, *p* = 0.7709. **(I)** Breakdown of spine morphology in juvenile mice. Statistical analysis by unpaired Student’s t-test for each spine class. For graphs D-F and I, data points represent individual animals, bars represent mean ± SEM.

We performed further analysis of spine morphologies and classified spines as mature or intermediate, to determine if there is an increase in mature spines. Protrusions with a mushroom-like morphology were classified as mature, and thin protrusions lacking a discernible spine head, or exhibiting a relatively dim, small head were identified as intermediate spines (**Figure 3 – figure supplement 1**). In P17-21 mice, WT and CKO mice showed similar proportions of mature spines (54% and 55% in WT and CKO, respectively, *n* = 4 mice per genotype; **Figure 3**), however mGFAP Smo CKO mice showed an apparent increase in protrusions with an intermediate morphology (16% and 26% in WT and CKO, respectively; **Figure 3**), though this was not statistically significant. This suggests that these intermediate protrusions reflect transitional morphologies as spines undergo maturation, and become more stable. Consistent with this, adult mGFAP Smo CKO mice trended toward a modest increase in the fraction of mature spines, compared to their WT controls (55% and 65% in WT and CKO, respectively, *n* = 4 mice per genotype; **Figure 3 – figure supplement 1**), whereas the fraction of intermediate spines was comparable at this age (27% and 23% in WT and CKO, respectively).

It is well established that the rate of synaptic turnover declines considerably as animals mature, reflecting an increase in stability of synaptic connections^26,27^. We next sought to investigate the long-term stability of individual spines in adult mice by imaging dendrites weekly for up to 6 weeks. We investigated the fraction of spines identified on the first day of imaging that persisted in subsequent imaging sessions using custom written MATLAB code (**Figure 3 – figure supplement 2**). We calculated the survival fraction by fitting to an exponential decay model. This analysis revealed a larger proportion of long-lived stable spines in mGFAP Smo CKO mice compared to WT control (71% and 79% in WT [*n* = 7 mice] and mGFAP Smo CKO [*n* = 5 mice], respectively, *p* = 0.0068, Extra sum-of-squares F test; **Figure 3**), suggesting an increase in stability of dendritic spines. Interestingly, among the population of dynamic spines, we found two distinct populations consisting of transient spines, which disappeared and did not reappear for the duration of the study, and recurrent spines, which disappeared and then subsequently reappeared. Our data showed that the shift towards increased spine stability in the mGFAP Smo CKO neurons was entirely due to a reduction in the proportion of recurrent spines, with nearly identical fractions of transient spines observed across both genotypes (**Figure 3**). These data suggest that structural plasticity is impaired in mGFAP Smo CKO mice. This deficit in structural plasticity emerges during the third week of postnatal development and persists into adulthood. Taken together, these data demonstrate that astrocyte-mediated Shh signaling is required for establishing and maintaining the organization of cell type specific cortical synapses.

### Shh signaling regulates Kir4.1 expression in cortical astrocytes

In both the developing and adult CNS, astrocytes directly eliminate synapses in an activity-dependent manner through MEGF10 and MERTK, two phagocytic receptors enriched in astrocytes^39^. To investigate whether these genes are negatively regulated in mGFAP Smo CKO mice, we examined their expression in the cortex of adult mice by quantitative PCR (qPCR). Expression of both *Mertk* and *Megf10* was comparable between WT and mGFAP Smo CKO mice (dCq values, *Mertk*: 5.97 ± 0.23 and 5.87 ± 0.06 in WT [*n* = 3 mice] and mGFAP Smo CKO [*n* = 6 mice], respectively, *p* = 0.54; *Megf10:* 8.02 ± 0.38 and 7.69 ± 0.18 in WT [*n* = 3 mice] and mGFAP Smo CKO [*n* = 6 mice], respectively, *p* = 0.39; **Figure 4 – figure supplement 1**). These data suggest that direct engulfment of synapses by astrocytes through these pathways is not regulated by Shh signaling and is not responsible for the increased spine density seen in mGFAP Smo CKO animals.

In the mature cerebellum, Shh signaling between Purkinje neurons and Bergmann glia regulates expression of the glial specific, inward-rectifying K^+^ channel, Kir4.1^21^. In the cortex, the distribution of Kir4.1 exhibits a laminar pattern similar to that of Gli1 astrocytes, such that expression is enriched in layers IV and Vb, and reduced in layer Va (**Figure 4**). High resolution, confocal analysis showed that Kir4.1 is localized in the processes of Gli1 astrocytes (**Figure 4**). Kir4.1 expression is localized to astrocytic endfeet and is found surrounding neuronal somata in the spinal cord and brain^40–42^. In WT mice, expression of Kir4.1 surrounded many NeuN-positive neuronal somata in layer V (**Figure 4**). In mGFAP Smo CKO mice, however, there was a pronounced reduction in the expression of Kir4.1 throughout the cortex, and peri-somal Kir4.1 expression was severely diminished (**Figure 4**). In order to quantify expression levels of Kir4.1, we measured transcript abundance in the cortex of mGFAP Smo CKO mice and WT controls by droplet digital PCR (ddPCR). In mGFAP Smo CKO mice, there was a 50% reduction in *Kir4.1* transcripts compared to WT controls (1.99 x 10^6^ ± 1.93 x 10^5^ copies/μl and 9.92 x 10^5^ ± 8.49 x 10^4^ copies/μl in WT [*n* = 6 mice] and mGFAP Smo CKO [*n* = 4 mice], respectively, *p* = 0.02). There was no difference in *Kir4.1* expression in CK2 Smo CKO mice compared to WT (1.88 x 10^6^ ± 3.77×10^5^ copies/μl, *p* = 0.95, *n* = 4 mice; **Figure 4**), suggesting that downregulation of *Kir4.1* is due to astrocyte-specific, and not neuronal, disruption of Shh signaling. Conversely, GLAST-CreER-mediated deletion of the Shh receptor *Ptch1*, a negative regulator of Shh signaling, dramatically upregulated Kir4.1 expression in the cortex (GLAST Ptch CKO; **Figure 4**). Kir4.1 expression is associated with glutamate uptake both *in vitro* and *in vivo*^43–45^. We therefore measured expression levels of the astrocyte-specific glutamate transporters *GLAST* and *GLT1*. There was no difference in expression of these genes in mGFAP Smo CKO mice compared to WT (*GLT1*: 1.08 x 10^7^ ± 6.6 x 10^5^ copies/μl and 9.58 x 10^6^ ± 2.1 x 10^5^ copies/μl in WT and mGFAP Smo CKO, respectively, *p* = 0.15, *n* = 3 mice per genotype; *GLAST:* 7.4 x 10^6^ ± 4.4 x 10^5^ copies/μl and 6.7 x 10^6^ ± 6.5 x 10^5^ copies/μl in WT and mGFAP Smo CKO, respectively, *p* = 0.41, *n* = 3 mice per genotype; **Figure 4 – figure supplement 1**). It should be noted that Kir4.1 was not restricted to Gli1 astrocytes in layers IV and Vb. Nevertheless, single transcript quantification of Kir4.1 transcripts in mGFAP Smo CKO show a pronounced reduction, suggesting that Kir4.1 expression in cortical astrocytes is regulated, in part, by Shh signaling.

**Figure 4:**
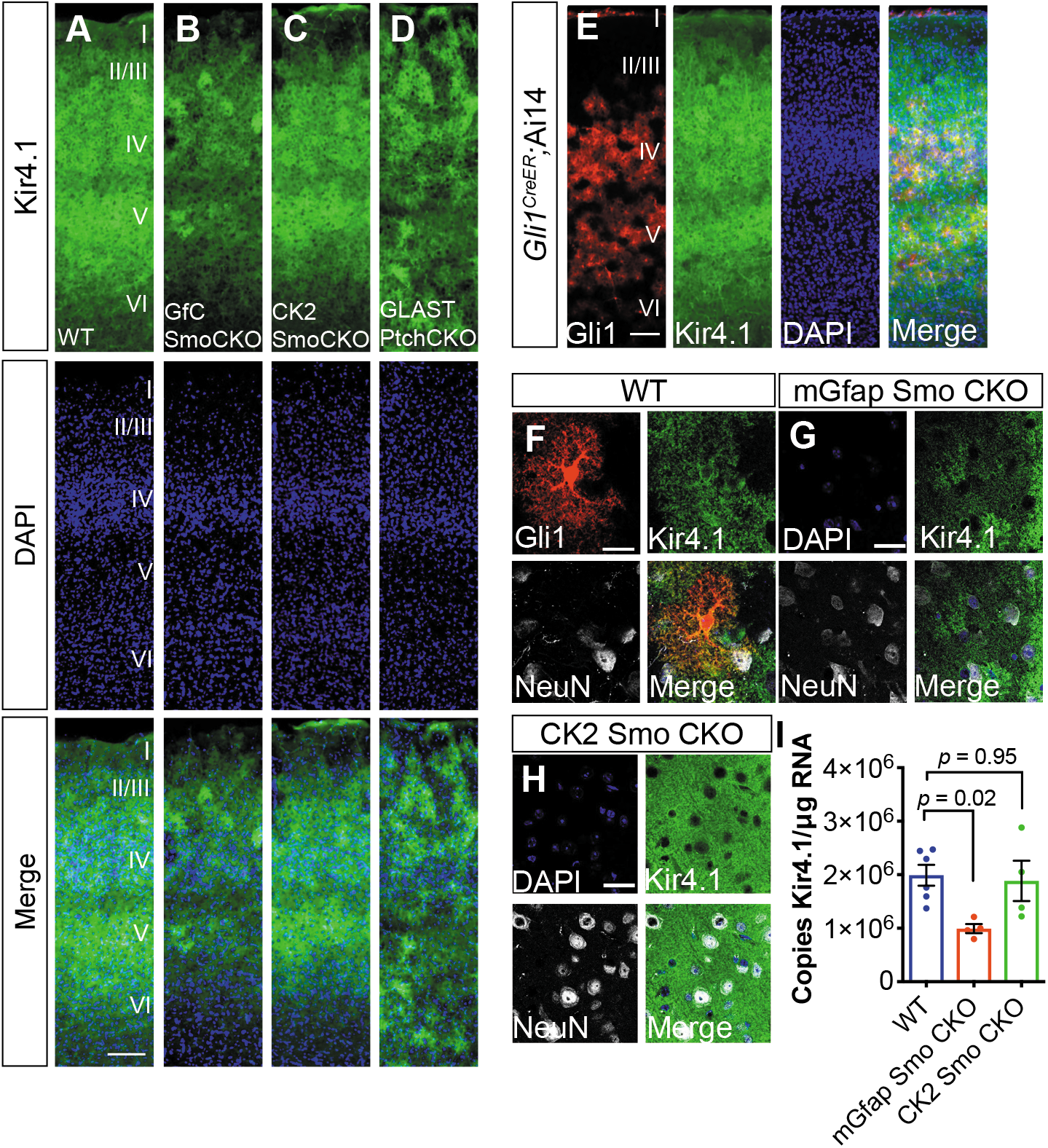
Shh signaling regulates Kir4.1 expression in the cortex. **(A-D)** Immunofluorescence for Kir4.1 in the cortex of adult WT (A), mGfap Smo CKO (B), CK2 Smo CKO (C) and GLAST Ptch CKO mice (D). Scale bar, 125 μm. **(E)** Fluorescence micrographs showing the distribution of Gli1 astrocytes (red) and Kir4.1 (green) in the cortex of an adult *Gli1Cre^ER^;*Ail4 mouse counterstained with DAPI (blue). **(F-G)** Confocal images of Gli1 (F, red) Kir4.1 (green) and NeuN (white) from adult WT (F), mGfap SmoCKO (G), or CamK2a SmoCKO (H) mice. Scale bar, 25 μm. (I) Gene expression levels in the cortex of Kir4.1 from WT (*n* = 6 animals) and mGfap Smo CKO or CK2 Smo CKO mice (*n* = 4 each). Data points represent individual animals and bar graphs represent mean value ± SEM. Statistical significance was assessed by one-way ANOVA with Tukey’s test for multiple comparisons. Significance stated on the graph is compared to WT

### Neurons exhibit hyperexcitability in mGFAP Smo CKO mice

To examine whether neurons in mGFAP Smo CKO mice exhibit disruptions in physiological activity, we performed whole-cell patch clamp recordings of layer V pyramidal neurons from coronal brain slices at P21, and examined action potential firing and membrane properties. Using current clamp, we recorded action potential spikes per injected current at multiple step currents from −300 pA to +650 pA. Our results revealed a significant increase in spike numbers with high current injections (>500 pA) and total overall spikes in mGFAP Smo CKO animals compared to WT control (326 ± 54 and 574 ± 71 spikes in WT [*n* = 11 cells from 4 animals] and mGFAP Smo CKO [*n* = 7 cells from 3 animals], respectively, *p* = 0.01; **Figure 5**). We also measured other membrane properties, including input resistance, resting membrane potential and tau, and found no significant difference between mGFAP Smo CKO and WT neurons (**Figure 5 – figure supplement 1**). However, we observed a significant decrease in action potential threshold (−30.6 ± 2.0 and −37.9 ± 2.0 mV in WT and mGFAP Smo CKO, respectively, *p* = 0.0441), accompanied by an increase in action potential amplitude (59.6 ± 2.6 and 73.3 ± 3.0 pA in WT and mGFAP Smo CKO, respectively, *p* = 0.0083), and a trending decrease in action potential ½ width (1.0 ± 0.92 and 0.76 ± 0.02 ms in WT and mGFAP Smo CKO, respectively, *p* = 0.05) in mGFAP Smo CKO neurons, consistent with an increase in neuron excitability (**Figure 5**). Together, these results suggest mGFAP Smo CKO neurons exhibit an increase in neuronal excitability consistent with excess extracellular K^+^.

**Figure 5:**
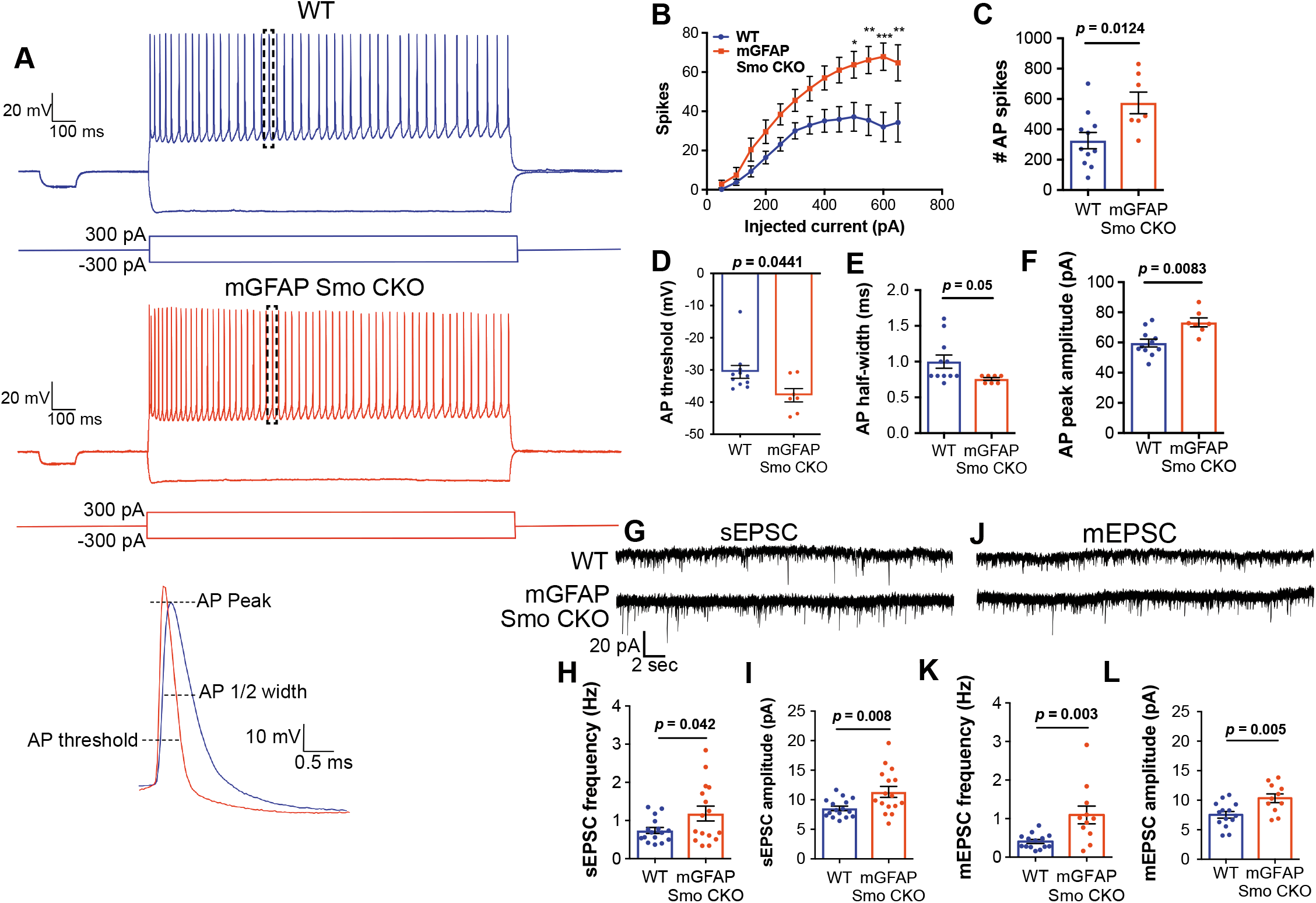
Neurons in mGFAP Smo CKO mice exhibit a significant increase in excitability and synaptic transmission. **(A)** Example traces of action potentials from layer V pyramidal neurons in WT and mGFAP Smo CKO mice at P21. Samples of action potential spikes (dashed lines, lower panel) describe AP threshold, AP peak amplitude, and AP ½ width of pyramidal neurons. **(B)** Line graph displays relationship between action potential spike numbers (y-axis) and current injection; mGFAP Smo CKO neurons exhibit an increase in AP spikes (>500 pA) compared to WT neurons. Statistical signifcance was assessed by two-way ANOVA with Sidak’s test for multiple comparisons and stated as mGFAP Smo CKO compared to WT at a given current (*n* = 11 and 7 cells from WT and mGFAP Smo CKO, respectively). **(C-F)** Bar graphs describe an increase in total AP spike numbers (C), reduction in AP threshold (mV) (D), reduction in AP 1/2 width (E), and increase in AP peak amplitude (pA) (F) in mGFAP Smo CKO neurons. **(G)** Example traces of sEPSCs from layer V pyramidal neurons recorded in the presence of picrotoxin from WT and mGFAP Smo CKO mice at P21. **(H-l)** Summary graphs of sEPSC frequency and amplitude (*n* = 14 and 11 cells from WT and mGFAP Smo CKO animals, respectively). **(J)** Example traces of mEPSCs from layer V pyramidal neurons recorded in the presence of picrotoxin and TTX. **(K-L)** Summary graphs of mEPSC frequency and amplitude (*n* = 16 cells per genotype). mGFAP Smo CKO neurons exhibit increases in both frequency and amplitude in sEPSCs and mEPSCs recordings. Statistical significance was assessed by unpaired Student’s t-test. For each graph, data points represent individual cells. All data are from at least 3 animals per genotype.

To examine excitatory synaptic transmission, we recorded spontaneous and miniature excitatory postsynaptic currents (sEPSCs and mEPSCs) in layer V pyramidal neurons. We found an increase in both the frequency (0.74 ± 0.08 and 1.18 ± 0.19 Hz in WT [*n* = 16 cells] and mGFAP Smo CKO [*n* = 16 cells], respectively, *p* = 0.042) and amplitude (8.54 ± 0.38 and 11.33 ± 0.91 pA in WT and mGFAP Smo CKO, respectively, *p* = 0.008) of sEPSCs in slices from mGFAP Smo CKO mice compared to WT control slices (4 animals per genotype; **Figure 5**). Furthermore, we observed an increase in the frequency (0.40 ± 0.05 and 1.10 ± 0.23 Hz in WT and mGFAP Smo CKO, respectively, *p* = 0.003) and amplitude (7.54 ± 0.57 and 10.34 ± 0.74 pA in WT and mGFAP Smo CKO, respectively, *p* = 0.005) in mEPSCs in the presence of tetrodotoxin (TTX), indicating an increase in postsynaptic response of neurons in mGFAP Smo CKO mice (*n* = 14 and 11 cells from WT and mGFAP Smo CKO, respectively, 4 animals per genotype; **Figure 5**). These data suggest that loss of Shh activity in astrocytes produces disturbances in both pre and postsynaptic activity. Moreover, these data suggest that astrocytes modulate both neuronal excitability and excitatory synaptic transmission in a Shh-dependent manner.

### Selective disruption of Shh signaling in astrocytes produces mild reactive gliosis

We previously demonstrated that cortical astrocytes in mGFAP Smo CKO mice upregulate GFAP expression and exhibit cellular hypertrophy, two classic hallmarks of reactive astrogliosis^17^. Astrocytes exhibiting these features were broadly distributed across cortical layers (**Figure 6**), in contrast to the distribution of Gli1 astrocytes which are found predominantly in layers IV and V. In addition to upregulation of GFAP, astrocytes in mGFAP Smo CKO mice show dramatic changes in morphological structure. Sholl analysis revealed no difference in the number of primary branches in mGFAP Smo CKO mice compared to WT controls (7.2 and 7.5 branches, WT [*n* = 9 cells] and mGFAP Smo CKO [*n* = 9 cells], respectively, 3 animals per genotype). However, astrocytes from mGFAP Smo CKO animals did show an increase in the number of higher order branches, the length of the longest tree (136.6 ± 17 and 212.1 ± 28 μm for WT and mGFAP Smo CKO, respectively, *p* = 0.035), and the total length of all processes (397 ± 66 and 674 ± 91 μm for WT and mGFAP Smo CKO, respectively, *p* = 0.025) compared to WT controls (**Figure 6**). Consistent with this, there was a significant increase in the number of branches intersecting concentric shells at various distances from the soma (**Figure 6**). Notably, in CK2 Smo CKO mice, GFAP staining in the cortex was indistinguishable from WT controls (**Figure 6 – figure supplement 1**), suggesting that astrocytes exhibit dramatic changes in morphological structure following astrocytic, but not neuronal, disruption of Shh signaling. Such changes in morphology are consistent with mild reactive gliosis, a complex cellular response of astrocytes to disturbances in physiological homeostasis^46^. Notably, the distribution of reactive astrocytes in mGFAP Smo CKO mice is broader than that defined by Gli1 expression, arguing against the idea that Shh signaling regulates the intrinsic state of astrocytes. Rather, reactive gliosis may instead reflect a cellular response to the mild, but persistent, aberrations in neuronal activity observed in mGFAP Smo CKO mice.

**Figure 6:**
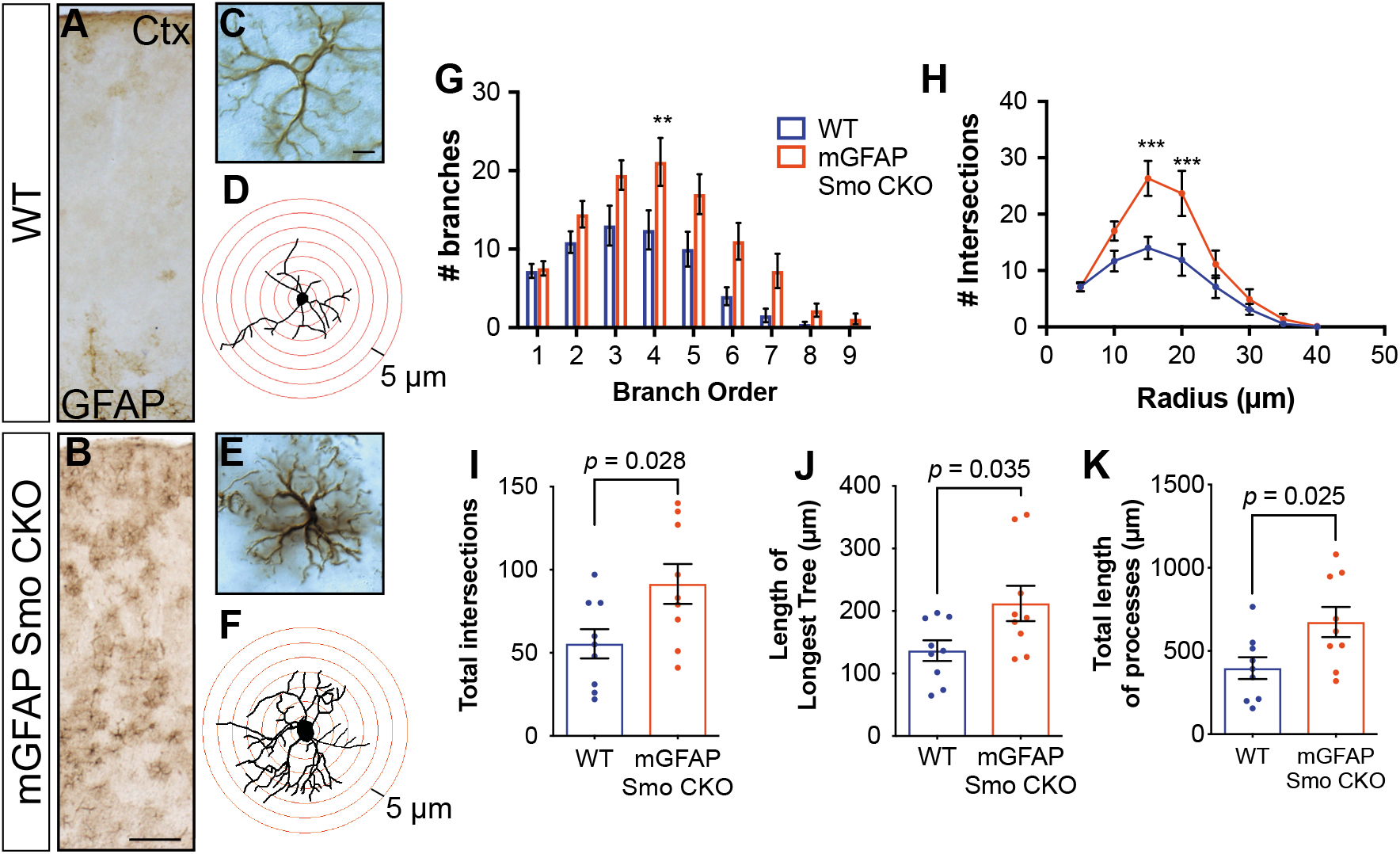
Disruption of Shh signaling in astrocytes results in widespread reactive changes in morphology. **(A-B)** Brightfield immunohistochemistry reveals a reactive upregulation of GFAP across cortical layers in mGFAP Smo CKO astrocytes. Scale bar, 125 μm. **(C & E)** Representative high-magnification images of both WT (C) and mGFAP Smo CKO (E) astrocytes are shown. Scale bar, 10 μm. **(D & F)** Representative traces of GFAP expression from WT (D) and mGFAP Smo CKO (F) cortical astrocytes. **(G & H)** Sholl analysis shows significant increases in complexity of mGFAP Smo CKO astrocytes compared to WT controls. Statistical significance was assessed by one-way ANOVA with Bonferroni’s for multiple comparisons. **(I-K)** Quantification of various morphological features of traced astrocytes. Statistical analysis performed by one-way ANOVA (G & H) and unpaired Student’s t-test (l-K). Data points represent individual cells. Graphs represent mean value ± SEM. *n* = 3 animals per genotype.

## DISCUSSION

The dynamic processes underlying organization of synapses during postnatal development are essential for establishing functional neural circuits. In this study, we demonstrate that astrocytes modulate refinement and reorganization of synaptic connectivity in the postnatal cortex in a Shh-dependent manner. Our data show that Gli1 astrocytes are enriched in deep cortical layers with a relative paucity in upper cortical layers. Selective disruption of Shh signaling in astrocytes leads to an overabundance of spines on the apical dendrites of layer V, but not layer II/III, cortical neurons. The overabundance of spines emerges during postnatal development, continues into adulthood and is accompanied by a reduction in dynamic turnover. mGFAP Smo CKO mice exhibit a pronounced reduction in Kir4.1 as well as an increase in neuronal excitability. Finally, we show that cortical astrocytes exhibit dramatic changes in morphology and GFAP expression, a phenotype consistent with mild reactive gliosis. Our data identify an essential role for astrocyte modulation of neuronal activity during postnatal development, facilitating activity-dependent reorganization of synaptic connectivity.

To accomplish selective disruption of Shh signaling in astrocytes, we used a Cre driver in which expression is regulated by the full-length mouse GFAP gene^22^. Single cell analysis in *mGFAP-Cre;Ai14* mice showed that nearly all cortical astrocytes undergo Cre-mediated recombination, while the fraction of recombined cortical neurons is less than 2%. Importantly, we observed recombined neurons predominantly in superficial layers, consistent with early Cre activity during late embryogenesis when layer II/III neurons are being generated. Notably, despite a small population of Smo null neurons in layer II/III, changes in spine density are not observed in these cells. In addition, a more complete deletion of Smo in excitatory neurons using the CamKIIαCre driver fails to alter spine density in layer V neurons, arguing against non-specific effects arising from the small population of recombined neurons in mGFAP Smo CKO mice. Importantly, Gli1 expression is nearly completely lost in mGFAP Smo CKO mice, with no loss in the number of total astrocytes^17^, demonstrating effective disruption of Shh activity in these mice. Because a small number of recombined oligodendrocytes were observed in mGFAPCre;Ai14 mice, the possibility that these cells may contribute to some of the observed phenotypes cannot be ruled out. However Gli1 expression is restricted predominantly to astrocytes in the adult forebrain^17^. Thus, any contribution from other cell types must be mediated by non-canonical, Gli-independent mechanisms.

In the postnatal cortex, Shh signaling from layer V neurons mediates synaptic connectivity with their layer II/III presynaptic partners^16^. Here, we show that, in addition to neuronal Shh signaling, Shh activity in astrocytes is required for the establishment and maintenance of cortical circuits. This is supported by two key observations in our study. First, we find that in mGFAP Smo CKO mice, apical dendrites of cortical neurons exhibit an increase in spine density that emerges during postnatal development and persists into adulthood. Importantly, genetic deletion of *Smo* in mature neurons instead of astrocytes failed to show any differences in spine density, pointing to a specific role for Shh signaling in astrocyte regulation of synaptic connectivity. Interestingly, deletion of Shh ligand using an EmxCre driver produces a reduction in spine density of basal dendrites^16^, whereas we found no difference in basal dendrite spine density, suggesting that Shh acts in distinct, cell type-specific ways to regulate synaptic connectivity of local cortical circuits. Indeed, whereas astrocytes express Gli1, neurons do not^17^, indicating differential transduction of Shh through canonical and non-canonical, Gli-independent pathways, respectively, in different cell types. Second, our data show fewer filopodia in juvenile mGFAP Smo CKO mice compared to WT controls along with a concomitant increase in spine stability. In the adult, spines continue to show deficits in dynamic turnover, demonstrating long-term impairments in structural plasticity. This suggests that Shh signaling is required not only for identifying appropriate synaptic partners during development, but also plays an important role in mediating synaptic plasticity.

In the postnatal and adult brain, we and others identified neurons as the source of Shh^16,17,21,47^. However non-neuronal sources, including astrocytes have been identified in the injured brain^48,49^. Beyond the CNS, epithelial cells have been reported as a source of Shh in the developing dentate gyrus^50^. It will be interesting to examine whether other non-neuronal sources are available to trigger astrocyte transduction of Shh.

We previously demonstrated that Gli1 expression is restricted to a subpopulation of cortical astrocytes^17^. Here, we extend these findings and demonstrate that the distribution of Gli1 astrocytes in the cortex is not uniform across cortical layers, but rather, occurs in a laminar-specific fashion. Cortical layers have been historically defined by neuronal populations and their specific functional properties and connectivity. However, emerging evidence suggests that astrocytes also exhibit cortical lamination patterns based on distinct gene expression profiles^51,52^. Our data show that Gli1 astrocytes are enriched in layers IV and V with a relatively sparse distribution in layer II/III. Interestingly, a recent study reported a population of astrocytes with a similar distribution that show enrichment of the Shh pathway^52^. These cells were also shown to regulate spine density of Layer V cortical neurons. It will be interesting to examine whether these cells correspond to Gli1 astrocytes reported in this study. Whereas astrocytes transducing Shh are enriched in deep cortical layers, the BMP antagonist, chordin-like 1 (Chrdl1) is preferentially enriched in upper layer cortical astrocytes^8^, suggesting that Gli1 and Chrdl1 may identify two distinct, but regionally complementary, astrocyte populations. Alternatively, Shh signaling may actively repress Chrdl1 expression in deep layer astrocytes. Notably, our observation that Gli1 astrocytes only comprise a fraction of astrocytes in layers IV and V suggests that, in addition to the laminar-specific distribution of these cells, there may be additional heterogeneity of astrocytes even within a given cortical layer. Whether and how the laminar distribution of astrocytes with distinct gene expression profiles confers functional specialization is not well understood and requires further study.

Interestingly, we observed that the disturbances in synaptic organization of cortical neurons in mGFAP Smo CKO mice occurs selectively in apical dendrites of layer V, but not layer II/III, neurons. Since apical dendrites of layer V neurons traverse through layer II/III, this suggests that Shh-dependent regulation of synaptic refinement is not mediated by direct interaction between Gli1 astrocytes and synapses. One possibility is that Gli1 astrocytes interact with neuronal soma in layer V and effectively modulate their excitability. Our observation that Kir4.1 expression in cortical astrocytes is regulated by Shh signaling suggests that mGFAP Smo CKO mice may experience deficits in local buffering of extracellular K^+^ leading to perturbations in neuronal activity. Indeed, a recent study demonstrated that Kir4.1 surrounds neuronal soma in the lateral habenula and regulates their firing properties^41^. Astrocytes play essential roles in buffering extracellular K^+^ through Kir4.1, and several studies demonstrate that loss of function of Kir4.1 impairs K^+^ uptake and increases neuronal excitability^44,45,53^. Alternatively, increased neuronal excitability may be due to disruptions in glutamate clearance. Although we did not observe changes in expression of either GLT1 or GLAST, deficits in Shh-dependent trafficking, localization or transporter function would lead to impairments in astrocyte-mediated glutamate clearance. Further work is needed to definitively identify the precise mechanism underlying the abnormal firing properties of neurons in mGFAP Smo CKO mice.

The establishment of appropriate synapse number and connectivity are tightly regulated processes that are profoundly shaped by experience or activity^34,54^. Our observation that mGFAP Smo CKO mice show increases in spontaneous neuronal activity and excitability, accompanied by a pronounced increase in spine density suggests that astrocytes contribute to activity-dependent processes that shape the organization of developing neural circuits in a Shh-dependent manner. The deficits in synaptic organization observed in mGFAP Smo CKO mice persist for the life of the animal. These deficits are not limited to synapse number but also include lower rates of spine turnover, suggesting long-lasting deficits in structural plasticity. Together with the observation that Shh guides local synaptic connectivity of cortical neurons^16^, these studies reveal that Shh signaling exerts considerable influence on the establishment and maintenance of cortical circuits.

## METHODS

### Key Resources Table

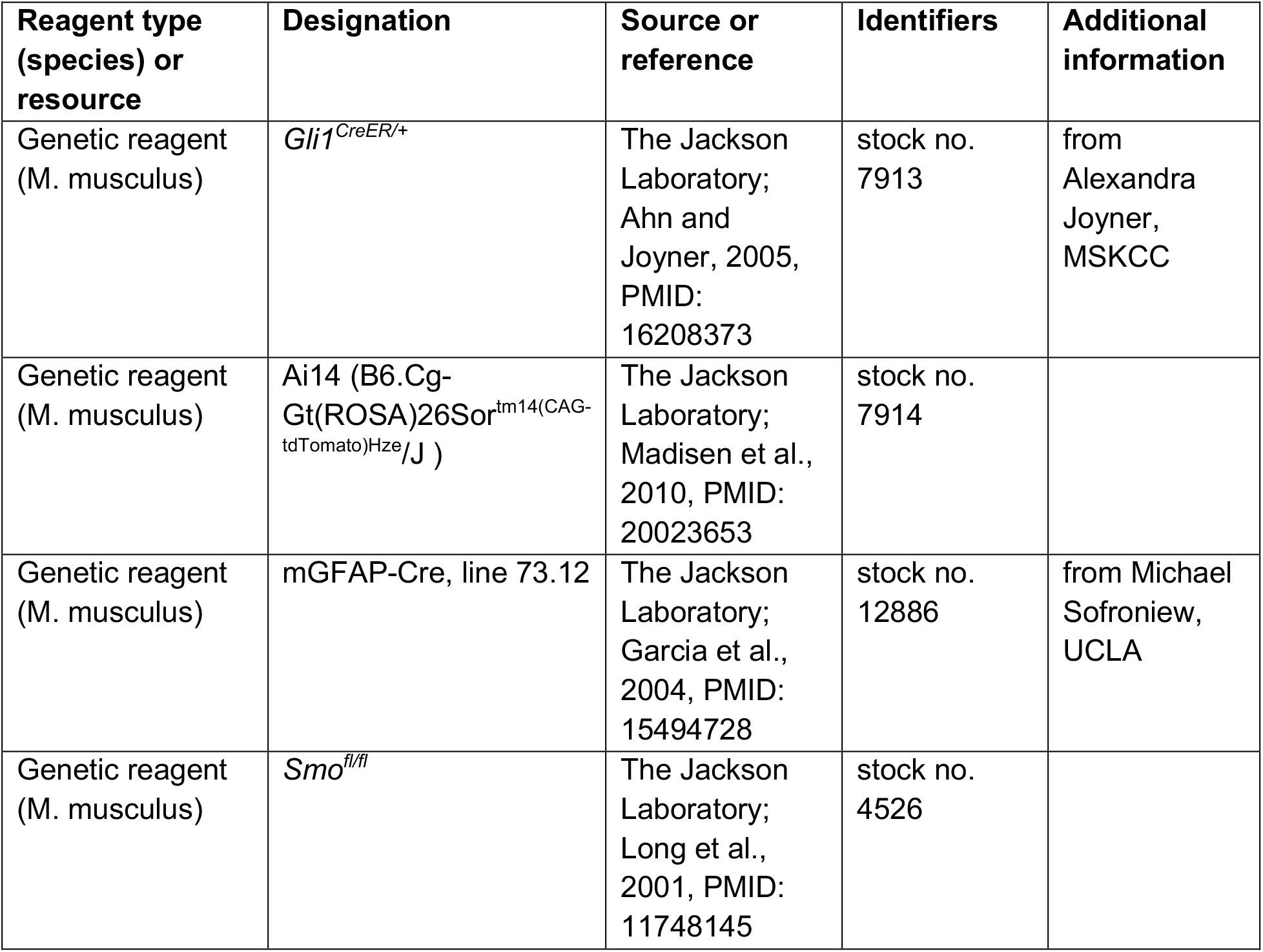
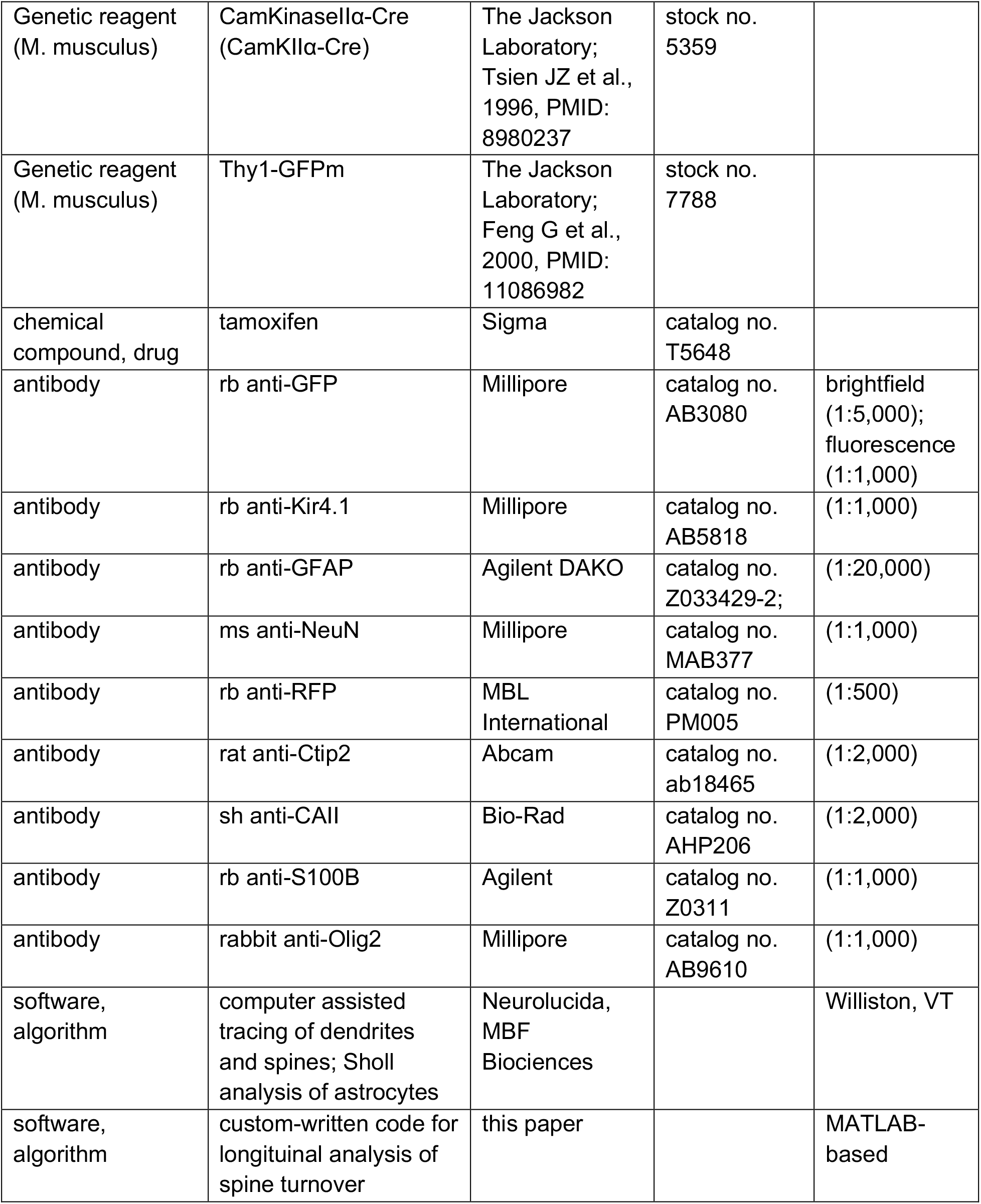

### Animals

All experiments were approved by Drexel University’s Institutional Animal Care and Use Committee and were conducted according to approved protocols. We used the following strains of transgenic mice on the C57BL/6 background: *Gli1*^*CreER*/+ 14^, Ai14 ^55^, Smo^*fl/fl* 56^, *mGFAP-Cre*^22^, *CaMKIIa-Cre*^57^ and *Thy1-GFPm*^25^. Mice of either sex were included in these studies.

### Tamoxifen

Tamoxifen was administered as previously described^17,20^. Briefly, tamoxifen (Sigma, T5648) was dissolved in corn oil to a final concentration of 20 mg/ml. Adult *Gli1*^*CreER*/+^*;Ai14* mice received 250mg/kg tamoxifen by oral gavage for three consecutive days, and tissues were analyzed two – three weeks later.

### Immunohistochemistry

Mice were deeply anesthetized by intraperitoneal injection of ketamine/xylazine/acepromazine. Animals were then transcardially injected with 100 units heparin and perfused with phosphate-buffered saline (PBS) followed by 4% paraformaldehyde (PFA, Sigma-Aldrich, St. Louis, MO). Brains were fixed overnight at 4°C and then transferred to 30% sucrose for at least 48 hours or until sectioned. Brains were cryosectioned (Leica CM3050S, Wetzlar, Germany) and collected in 40 μm sections. Sections were stored in 0.1M Tris-buffered saline (TBS) with 0.05% sodium azide at 4°C. Immunohistochemistry was performed using the following primary antibodies: rabbit anti-GFP (AB3080; Millipore, Burlington, MA), rabbit anti-Kir4.1 (AB5818; Millipore), rabbit anti-GFAP (Z033429-2; Agilent DAKO, Santa Clara, CA), mouse anti-NeuN (MAB377; Millipore), rabbit anti-RFP (PM005; MBL International, Woburn, MA), rat anti-Ctip2 (ab18465; Abcam, Cambridge, UK), sheep anti-CAII (AHP206; Bio-Rad), rabbit anti-S100β (Z0311, Agilent), and rabbit anti-Olig2 (AB9610; Millipore). Sections were washed three times in 0.1M TBS at room temperature (10 minutes/wash). Sections were then blocked in 10% Normal Serum with 0.5% Triton-X (Sigma-Aldrich) for one hour at room temperature and incubated with primary antibody in 0.5% Triton-X at 4°C overnight. For fluorescent immunohistochemistry, sections were rinsed the following day in 0.1M TBS and incubated in Alexafluor-conjugated secondary antibodies with 10% Normal Serum and 0.1M TBS and incubated at room temperature for two hours. The sections were then rinsed in 0.1 TBS and incubated in DAPI (1:50,000; Life Technologies, Carlsbad, CA) for 15 minutes. Sections were rinsed again in 0.1M TBS and mounted onto microscope slides (Fisherbrand, Waltham, MA) and coverslipped using ProLong Gold Antifade Mountant (Invitrogen, Carlsbad, CA) and Fisherfinest Premium Cover Glass. For brightfield immunohistochemistry, sections were rinsed as previously described but were incubated with biotinylated secondary antibodies against the primary antibody species (Vector Laboratories, Burlingame, CA) for one hour. Sections were then placed in Avitin-Biotin Complex solution (Vector) for one hour and visualized using 3,3’-diaminobenzidine (DAB Peroxidase Substrate Kit, Vector). Sections were then mounted onto slides and coverslipped with DPX Mountant (Fisher).

### Quantification of dendritic spine density

Analysis of spine density was performed in a blinded study design. WT controls were derived from Cre-negative littermates. Neurons were traced using Neurolucida (MicroBrightField Biosciences, Williston, VT) and individual spines were marked. Apical dendrites of layer V neurons were counted from ~100 μm below the primary bifurcation through the apical tuft. Apical dendrites of layer II/III cells were analyzed just below the primary bifurcation through the apical tuft. Basal dendrites were analyzed beginning ~50 μm from the soma through the end of the processes. CA1 hippocampal neurons were analyzed from just below the primary bifurcation through the apical tuft. All analysis was performed on an upright Zeiss microscope using a 63X oil objective.

### Cranial window

Cranial window surgeries were adapted from a published protocol^35^. Immediately prior to each surgery, mice received a subcutaneous injection of carprofen (Rimadyl, 5 mg/kg), and a follow-up injection was given 24 hours after surgery. Mice were deeply anesthetized with isoflurane (5%) in an induction chamber, and then transferred to a stereotaxic apparatus where they inhaled 1.5% isoflurane continuously for the duration of the surgery. The scalp was then removed and a metal bar (~1 cm long) with threadings for screws was affixed to the skull, using superglue and dental acrylic to form a head post that sealed off the skin while leaving the right parietal bone exposed. After allowing 24 hours for the glue and acrylic to set, the mouse was returned to the stereotaxic apparatus and head-fixed using the headpost. A circular craniotomy ~3 mm in diameter and centered 2.5 mm lateral and posterior to bregma was performed. The exposed dura was treated with saline-soaked gel foam until any minor bleeding ceased. Finally, a 3 mm glass coverslip was gently pressed onto the dura and sealed in place using superglue and dental acrylic. Adult mice were allowed at least 1 week for recovery. To mitigate the tendency of rapid bone growth to destabilize the windows at younger ages, juvenile mice were allowed 24 hours to recover.

### Two-photon imaging

A commercial two-photon microscope (Bruker, Billerica, MA) with a tunable Ti:Sapphire laser (Coherent, Santa Clara, CA) was used for all experiments. The laser was set to 920 nm and power was tuned on the sample as necessary to obtain consistently high-quality images without damaging the tissue. Mice were head-fixed and anesthetized with continuous isoflurane (1-1.5%) during imaging. Under a 20X water immersion objective, GFP-expressing neurons were identified visually and then imaged down to the cell body to determine layer. From the apical dendrite tuft, individual dendrite segments that projected primarily along the xy-plane were selected for analysis, and z-stacks (0.5 μm step size) were collected. To track individual dendrites, repeated imaging was generally performed on the following schedule (in days, relative to first imaging session): 0, 1, 2, 7, 14, 21, 28, 42. Some data from studies using other schedules were also included. For juvenile mice, this schedule was: 0, 1, 2.

### Analysis of spine dynamics

Every effort was made to use Cre-negative, littermate controls for these experiments. However, it was not always possible to match littermate controls in the final data sets presented here, either due to the absence of the GFP allele in the Cre-negative controls, or due to the loss of head caps securing the cranial windows, over time. However, it should be noted that all data presented here represent Cre-negative controls from the appropriate strain, if not necessarily from the same litter.

To analyze spine turnover, we performed side-by-side manual comparison of dendritic protrusions using ImageJ. Z-stacks from two time points were analyzed, and individual protrusions were identified as present or absent in each stack. For each mouse, a minimum of 150 protrusions were analyzed, although the average number of protrusions analyzed per mouse was 250-300. These comparisons were conducted by trained observers blinded to mouse genotype. Long, thin protrusions lacking a bulbous head were identified as filopodia, and any other prominent dendritic protrusion that extended > 0.4 μm from the shaft was counted as a spine. The filopodial fraction was calculated as the proportion of all analyzed features present at the first imaging timepoint that were identified as filopodia. The position and dynamic status (stable/eliminated/formed) of each spine was recorded using ImageJ’s CellCounter plugin. Each neuron’s turnover ratio was calculated as:

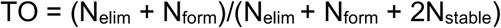

For longitudinal analysis of spine lifetime, the individual features identified in our manual ImageJ analysis were then imported into custom written MATLAB (Mathworks, Natick, MA) code and manually tracked across all imaged time-points using custom-written software. This code facilitated the efficient manual tracking of each feature by iteratively estimating the z-plane, within each stack, in which that feature would be expected to appear, assuming a linear relationship. Using a graphical user interface, the user was presented with a series of frames centered on that plane and allowed to make a determination of the feature’s presence or absence simply by left-clicking on the feature, or in the case of absence right-clicking on the location it otherwise would have been, using the best frame presented (**Figure 3 – figure supplement 2**). Due to the iterative nature of this procedure, each dataset was run through at least twice to ensure that not only were the determinations of presence/absence accurate, but that the marked locations of every feature were optimal.

Spines that were observed consistently across all time points were classified as stable. Spines that disappeared at some time point and were observed again in a subsequent time point were classified as recurrent. Spines that disappeared and never reappeared were classified as transient. To compare long-term dynamics between WT and mGFAP Smo CKO mice, we examined the survival curves, S(t), for each mouse, where:

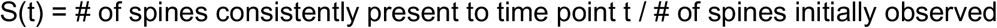

Survival curves for each mouse were pooled by genotype and fit to a single-phase exponential decay model:

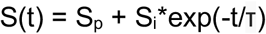

where S_p_ and S_i_ represent the relative fractions of permanent and impermanent spines respectively and τ is the characteristic lifespan of impermanent spines. Fitting and statistical comparison of survival data was performed with PRISM 6 (GraphPad, San Diego, CA) software. To analyze morphology, we examined data from the first imaged day, and classified individual protrusions as filopodia, intermediate, or mushroom spines. For each protrusion, a mean projection image was generated, averaging the frame on which the protrusion was manually marked in the original analysis and the 2 neighboring frames (1 above and 1 below, effectively averaging over 1.5 um of depth in the z axis). Each image was cropped, centering around the protrusion’s XY coordinates and a marker was drawn on the image to indicate which protrusion was under consideration. These images were then presented to the scorer sequentially for scoring. Protrusions exhibiting a bright, bulbous head were scored as mushroom. Thin protrusions exhibiting a relatively dim, small head, or no head, were scored as intermediate. Dim, elongated protrusions that lacked any head were scored as filopodia.

### Transcript Quantification

Mice were deeply anesthetized by intraperitoneal injection of ketamine/xylazine/acepromezine before rapid decapitation. Cortices were dissected into TRIzol reagent (Thermo-Fisher, Waltham, MA) and RNA was extracted according to the standard protocol. RNA was then purified with an RNeasy Micro Kit (Qiagen, Hilden, Germany) and subsequently reverse transcribed to cDNA using the High-Capacity cDNA Reverse Transcription Kit (Thermo-Fisher). Droplet digital PCR (ddPCR) was performed with the QX200 Droplet Digital PCR System using Evagreen Supermix (Bio-Rad, Hercules, CA). Alternatively, for *Megf10* and *Mertk*, qPCR was performed with the CFX96 Touch Real-Time PCR Detection System (Bio-Rad) using PowerUp SYBR Green Master Mix (Thermo-Fisher). PrimePCR ddPCR Expression EvaGreen Assay (Bio-Rad) was used for *Gapdh* primers; all other primers were designed using NCBI Primer-BLAST as follows: *Kir4.1* F, GCTGCCCCGCGATTTATCAG; *Kir4.1* R, AGCGACCGACGTCATCTTGG; *GLAST* F, TCCTCTACTTCCTGGTAACCC; *GLAST* R, TCCACACCATTGTTCTCTTCC; *Glt1* F, CATCAACAGAGGGTGCCAAC; *Glt1* R, CACACTGCTCCCAGGATGAC; *Megf10* F, CTCACTGCTCTGTCACTGGGTG; *Megf10* R, GGTAGCTGATTCTGTGCCGTGT; *Mertk* F, AAACTGCATGTTGCGGGATGAC; *Mertk* R, TCCCACATGGTCACGCCAAA. ddPCR absolute quantification of targets was calculated using Quantasoft Version 1.7 software (Bio-Rad). Samples were run in triplicate, and *Gapdh* was quantified for each sample as a loading control with no difference detected between sample groups.

### Slice electrophysiology

Mice aged P21 were anesthetized with intraperitoneal injection of euthasol (0.2 ml/kg) and brains were rapidly removed. 300 μm coronal slices were cut using a vibratome tissue slicer (Leica) and transferred to a holding chamber, submerged in oxygenated artificial cerebrospinal fluid (ACSF, in mM: 124 NaCl, 2.5 KCl, 1.25 NaH_2_PO_4_, 2 CaCl_2_, 1 MgSO_4_, 26 NaHCO_3_, and 10 dextrose, pH 7.4) at 36°C for 1 hour and then maintained at room temperature. Individual slices containing the barrel cortex were placed into a recording chamber immersed in oxygenated ACSF mounted on an Olympus upright microscope (BX51). Neurons were visualized with infrared differential interference video microscopy.

Whole-cell current clamp was used to record action potentials from layer V pyramidal cells using patch electrodes with an open tip resistance of 7–10 MΩ. Patch electrodes were filled with potassium gluconate internal solution (in mM): 120 potassium gluconate, 20 KCl, 4 ATP-Na, 0.3 Na_2_GTP, 5 Na-phosphocreatine, 0.1 EGTA, 10 HEPES, pH 7.3, 305 mosmol/l) and action potential responses were measured in response to various step currents from −300 pA to +650 pA with 50 pA increments. Action potentials were described as spike numbers per depolarized current injections. The resting membrane potential, input resistance, action potential (AP) threshold, AP half-width, and peak AP amplitude were also measured.

Whole-cell voltage-clamp recordings were obtained from layer V pyramidal cells using patch electrodes with Cs^+^-containing solution (in mM): 120 Cs-gluconate, 5 lidocaine, 6 CaCl_2_, 1 Na_2_ATP, 0.3 Na_2_GTP, and 10 HEPES (pH 7.3 adjusted by CsOH). To record spontaneous excitatory postsynaptic currents (sEPSCs), cells were held at a membrane potential of −70mV in the presence of GABA_A_ receptor antagonist picrotoxin (PTX; 100 μM, Sigma-Aldrich) and recorded for 5 minutes. Miniature EPSCs (mEPSCs) were then recorded for an additional 5 minutes in the presence of both picrotoxin and tetrodotoxin (TTX, 0.5 μM, Hello Bio, Princeton, NJ). All recordings were conducted with Axon MultiClamp 700B amplifier (Molecular Devices, San Jose, CA). The sEPSCs and mEPSCs frequency and amplitudes were measured by averaging 5 sweeps from the onset of recording with Clampfit 9.2 software (Molecular Devices). The s/mEPSCs were detected and characterized using a sample template for the 5-minute data period. Based upon the selected sample template, the frequency (number of event detections) and amplitude of events were measured using a threshold set in Clampfit.

### Statistical Analyses

Statistical analyses used for various datasets are indicated both in the figure legends and in the text. Prism 6 software (GraphPad) was used for all analyses and to generate graphs.

## Acknowledgements

We would like to thank Pooja Sakthivel and Nikhil Karmacharya for technical assistance and Dr. Monica Truelove-Hill for assistance with statistical analyses. We also thank Dr. Michael Akins for helpful discussions.

---

**Figure 2 – figure supplement 1:**
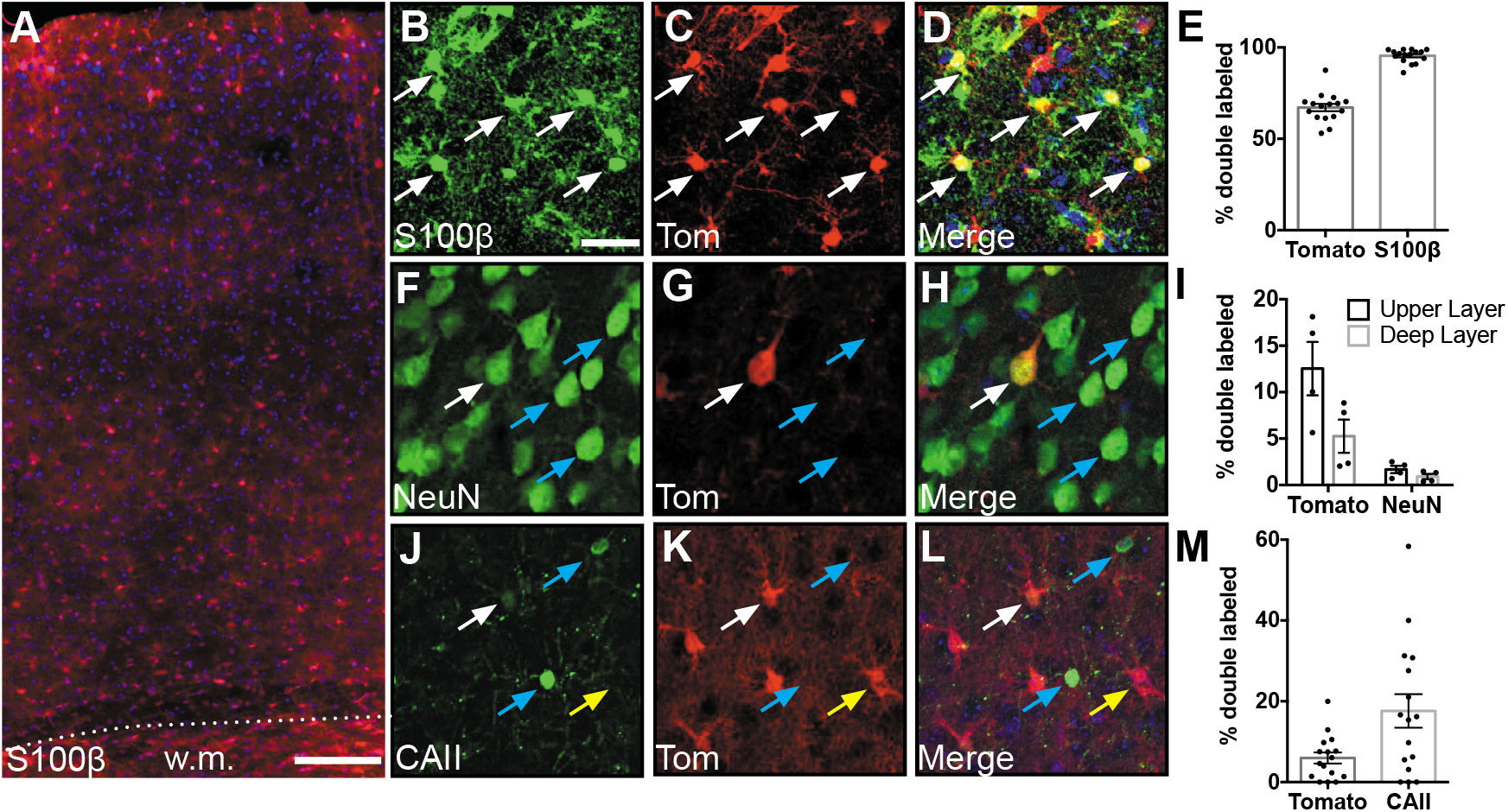
mGfap-Cre recombination occurs primarily in astrocytes. **(A)** Representative image of tdTomato expression in the cortex of *mGfap-Cre;*/Ai14 mice, w.m., white matter. Scale bar, 125 μm. **(B-D)** Expression of the astrocyte marker S100β and tdTomato in somatosensory cortex. Scale bar, 25 μm. **(E)** Fraction of recombined cells identified as astrocytes in the cortex of adult *mGfap-Cre*;/Ai14 mice. **(F-H)** Expression of the neuronal marker NeuN and tdTomato in somatosensory cortex. **(I)** Fraction of recombined cells identified as neurons. **(J-L)** Expression of the oligodendrocyte marker CAII and tdTomato in somatosensory cortex. **(M)** Fraction of recombined cells identified as oligodendrocytes. White arrows show double-labeled cells, blue arrows show cells that express the indicated marker but are tomato-negative, and yellow arrows show single-labeled tomato-positive cells. Data points represent individual brain sections analyzed. Bar graphs represent mean ± SEM. *n* = 2 animals, four sections per animal.

**Figure 2-figure supplement 2:**
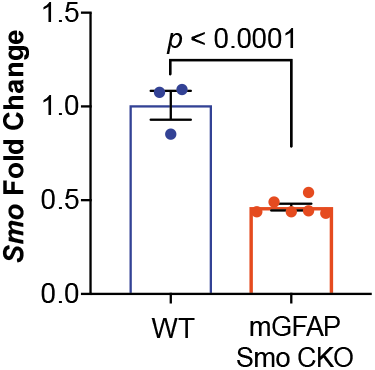
mGFAP Smo CKO mice have reduced levels of *Smo*. Quantitative PCR of cortical RNA shows a significant reduction in *Smo* expression. Data points represent individual animals, bars represent mean ± SEM. Statistical analysis assessed by unpaired Student’s t-test.

**Figure 2 – figure supplement 3:**
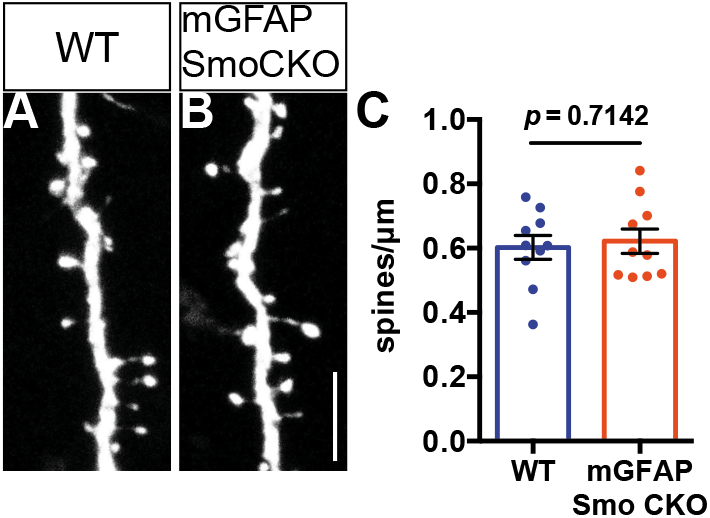
Spine density on layer V basal dendrites are not affected in adult mGFAP Smo CKO mice. **(A-B)** Representative images of basal dendrites analyzed from layer V neurons expressing *Thy1GFPm* in both WT (*Smo^fl/fl^*) and mGFAP Smo CKO (*mGfap-Smo^fl/fl^*) animals. Scale bar, 5 μm. **(C)** Graph comparing the density of spines on basal dendrites from WT and mGFAP Smo CKO animals. Data points represent individual cells, bars represent mean ± SEM. Statistical significance was assessed using unpaired Student’s t-test. *n* = 3 animals per genotype.

**Figure 3 – Supplement 1.**
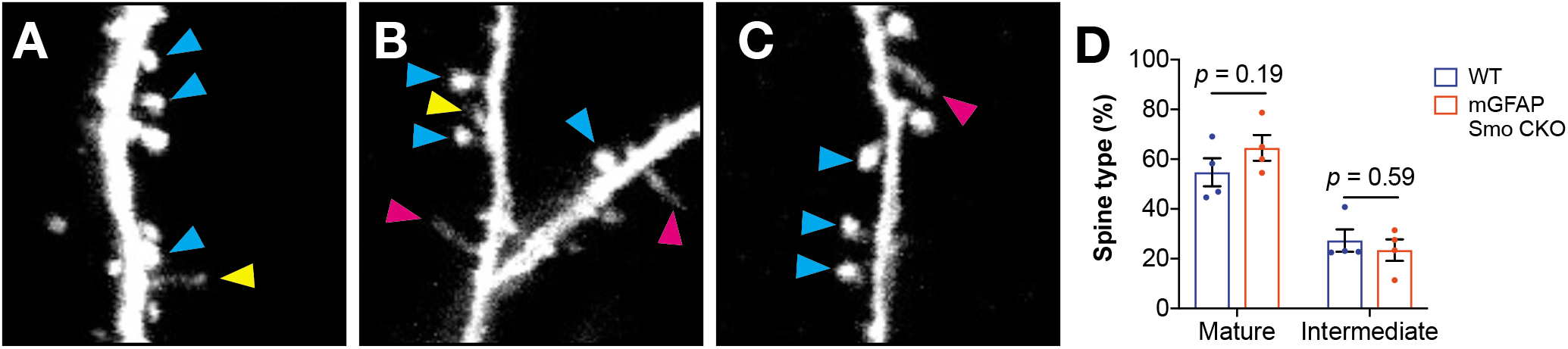
**(A-C)** Several examples of protrusions classified as mushroom (blue arrowheads) or intermediate (yellow arrowheads) spines, and filopodia (pink arrowheads). Note the thin stalk and small bulbous head of the intermediate spine in A, whereas filopodia lack a discerible head (B and C). **(D)** The fraction of protrusions classified as mushroom or intermediate in adult mGFAP Smo mice and WT controls. Data points represent individual animals, bars represent mean ± SEM. Statistical analysis by unpaired Student’s t-test.

**Figure 3 – Supplement 2.**
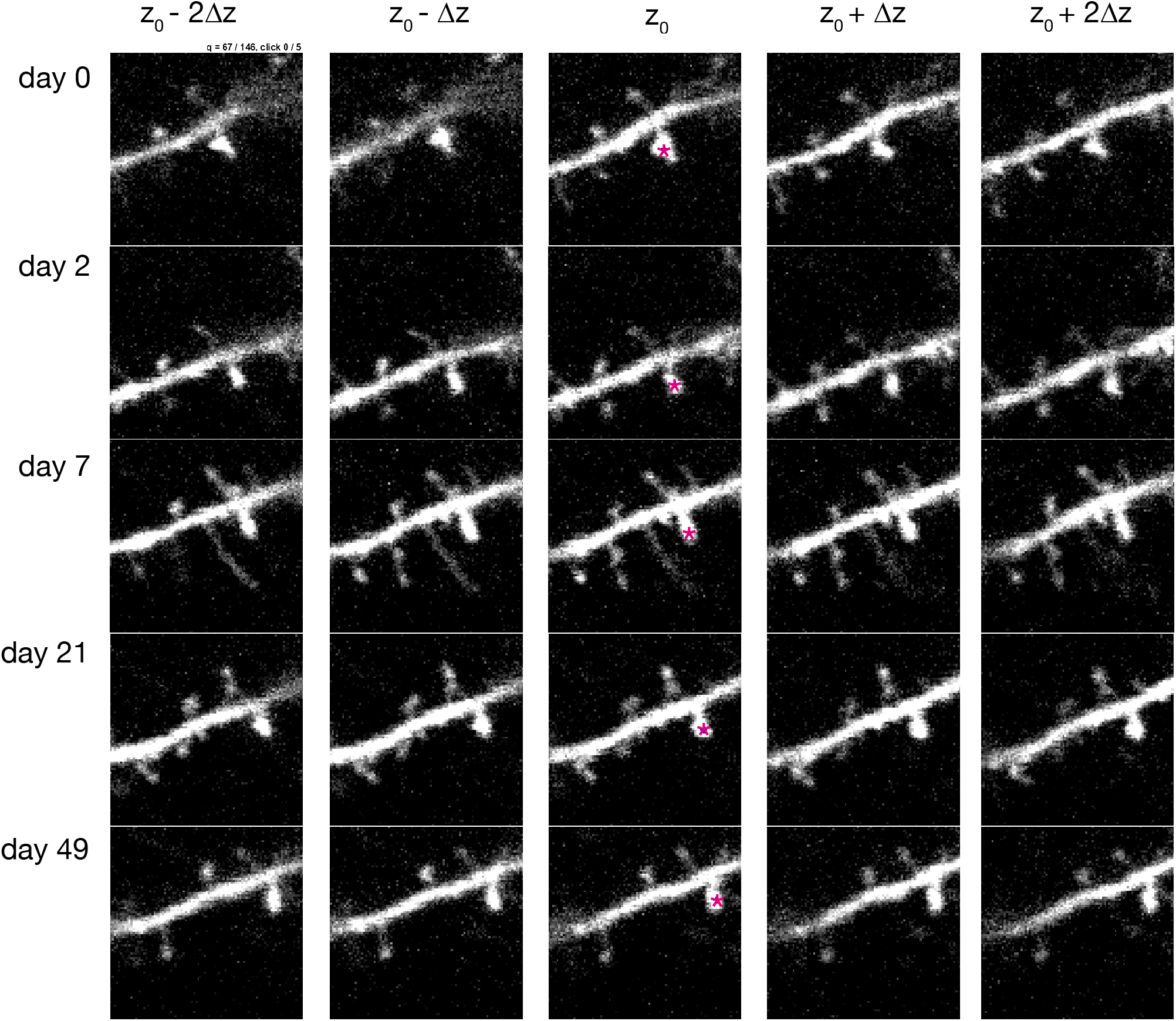
mGfap-Cre recombination occurs primarily in astrocytes. Screenshot depicting the graphical user interface used for longitudinal tracking of spines across multiple days of imaging. A spine identified in the first imaging session (pink star, top central panel) is presented to the user for scoring. The same spine is manually tracked across subsequent days (rows). For each day, multiple frames above and below the estimated z-plane corresponding to that spine, Z0, are also presented (columns). For each timepoint, the user then clicks on the spine’s head using the frame that best captures the spine. In this case, the frame jump, Δz, was set to 1, and 4 neighboring frames are presented; these parameters can be adjusted as necessary.

**Figure 4 – figure supplement 1:**
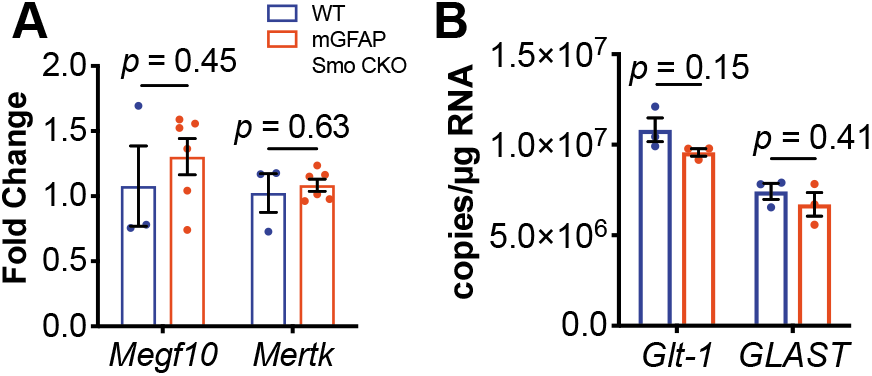
Quantititive PCR analysis of known spine and synapse modulators show no change in expression in mGFAP Smo CKO mice. **(A)** Quantitive PCR shows no change in *Megf10* or *Mertk* in mGFAP Smo CKO animals compared to controls (*n* = 3 and 6 samples of cortical RNA from WT and mGFAP Smo CKO animals, respectively). Statitistical significance assessed by unpaired Student’s t-test of the quantification cycles for each sample. Error bars represent standard error of the mean. **(B)** Droplet digital PCR shows no difference in the number of trancripts of either *GLT-1* or *GLAST* in mGFAP Smo CKO animals (*n* = 3 cortical RNA samples per genotype). Statistical significance assessed by unpaired Student’s t-test of the copy number for each target in each genotype. Error bars represent standard error of the mean.

**Figure 5 – figure supplement 1.**
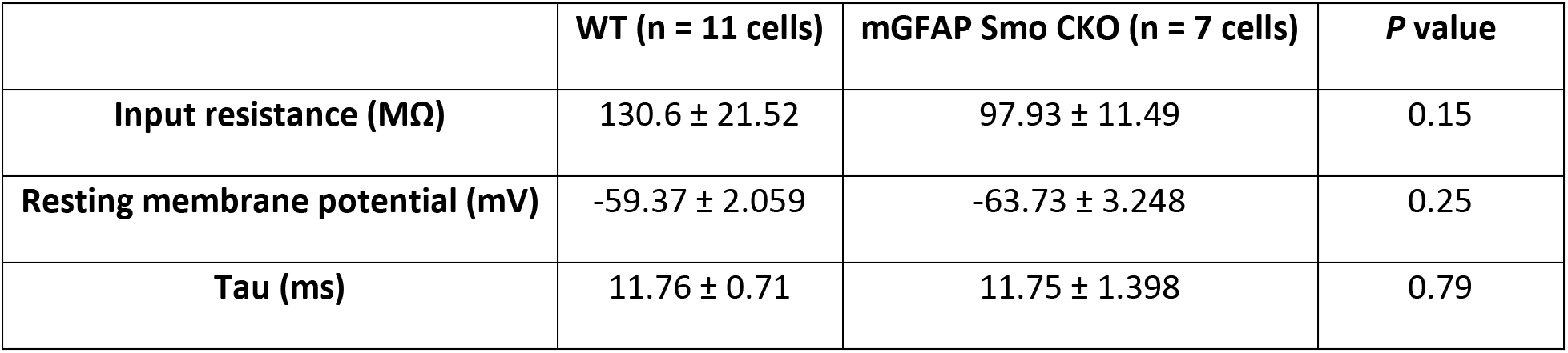
mGfap-Cre recombination occurs primarily in astrocytes. Membrane properties of mGFAP Smo CKO layer V pyramidal neurons. There is a significant decrease in input resistance (MΩ) in mGFAP Smo CKO neurons compared to WT neurons. No significant differences were observed in resting membrane potential (mV), rheobase (pA), and tau (ms) in mGFAP Smo CKO neurons. Statistical significance was assessed by Mann-Whitney U test (input resistance and tau) and unpaired Student’s t-test (resting membrane potential).

**Figure 6 – figure supplement 1:**
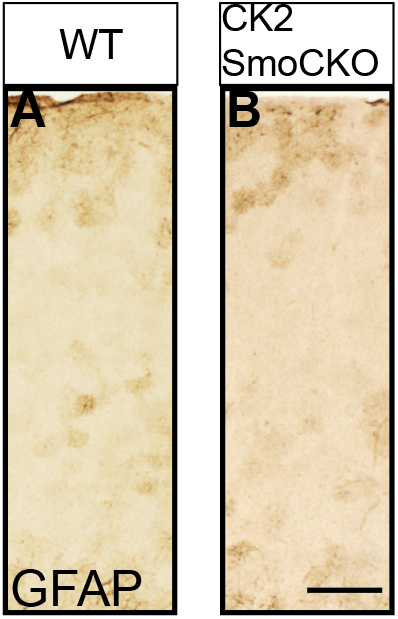
Reactive gliosis does not occur when neuronal Shh signaling is disrupted. Brightfield immunohistochemistry for GFAP in the cortex of an adult *CamKlla;Smo^fl/fl^* animal (B) and littermate control (A) shows comparable levels of GFAP expression. Scale bar, 125 μm.

